# Structural basis of metalloid transport by the arsenite efflux pump ArsB

**DOI:** 10.64898/2026.02.19.706881

**Authors:** Shivansh Mahajan, Kemal Demirer, William M. Clemons, Douglas C. Rees

## Abstract

Bacteria resist toxic arsenite (As^III^) in their environments by actively pumping the metalloid out of the cell via efflux pumps such as ArsB. However, the mechanism of extrusion remains poorly understood, which hinders the development of engineered bioremediation strategies. We report high-resolution cryo-EM structures of ArsB from the arsenic-tolerant bacterium *Leptospirillum ferriphilum*. ArsB adopts an inverted two-fold repeat architecture resembling that of other ion transporter (IT) superfamily proteins. Structures determined in the presence of As^III^ and antimonite (Sb^III^) reveal that the metalloid substrates interact with polar residues at the core of the transmembrane domain primarily via hydrogen bonding. Mutagenesis and *in vivo* functional assays support these interactions. Our ArsB structures represent an ‘inward-facing’ conformation, where the metalloid-binding site is exposed to the cytoplasm, suitable for metalloid capture. Furthermore, we demonstrate that As^III^ resistance conferred by ArsB varies with external pH, supporting that ArsB is a proton (H^+^)-coupled secondary transporter. Mutagenesis, *in vivo* functional assays, and pK_a_ estimation imply that conserved aspartate residues near the metalloid-binding site likely mediate the H^+^-coupling mechanism. Our findings provide structural insights into metalloid recognition and H^+^/metalloid antiport in ArsB, laying a foundation for further elucidation of the molecular basis of toxic metalloid detoxification in bacteria.

## Introduction

Arsenic is an environmental toxin that contaminates groundwater and crop fields in many regions across the globe, posing widespread public health risks^1,2^. Many bacteria and some archaea detoxify arsenic and other toxic metalloids, such as antimony, via metalloid-specific efflux pumps^3^. This is especially critical for bacteria that thrive in arsenic-rich environments, such as *Leptospirillum ferriphilum*, which is found in acid-mine drainage and hot springs, and can tolerate up to 60 mM arsenite^4^. The most widely recognized of these efflux pumps is ArsB, an integral membrane transporter encoded within the *ars* operon^5,6^. ArsB keeps the intracellular concentrations of the metalloid at subtoxic levels by capturing and exporting trivalent arsenite (As^III^) or antimonite (Sb^III^). This export is enacted either by energetically coupling metalloid efflux with the proton motive force (PMF) across the membrane (secondary active transport) or by association of ArsB with the cytoplasmic ATP hydrolase ArsA, which improves efflux efficiency by coupling to ATP hydrolysis (primary active transport)^7,8^. These distinct modes of metalloid export confer tolerance to a wide range of metalloid concentrations. The underlying molecular mechanism of ArsB is of interest for understanding heavy metal detoxification in living systems and can be leveraged to engineer robust, sustainable arsenic bioremediation strategies^9,10^.

ArsB is a member of the ion transporter (IT) superfamily that includes highly versatile secondary transporters that translocate charged substrates across the cell membrane^11,12^. Other members of this superfamily have been structurally and mechanistically characterized, including the divalent anion-sodium symporters (DASS)^13–16^, the p-aminobenzoyl glutamate transporters^17^, and the tripartite ATP-independent periplasmic (TRAP) transporters^18,19^. These transporters bind and transport a broad range of ‘oxyanionic’ substrates, such as di/tricarboxylate intermediates of the tricarboxylic acid cycle, phosphate, and sialic acids^13,16,20,21^. Despite substantial sequence homology, ArsB is proposed to be mechanistically distinct from other IT proteins for three key reasons. First, As^III^ exists as a neutral trihydroxylated species [AsOH)_3_] at physiologically-relevant pH^22^, in contrast to the negatively charged substrates transported by other IT proteins. Second, while most well-characterized IT proteins energetically couple substrate transport to the Na^+^-gradient across the membrane, ArsB couples metalloid transport to the proton (H^+^)-gradient across the membrane^18,23^. Third, IT proteins are typically ‘Na^+^/substrate symporters’, whereas ArsB is a ‘H^+^/metalloid antiporter’^8^. For DASS family transporters, the mechanisms of both substrate binding and Na^+^-coupled symport are well characterized at the structural level^13^. However, no structural information is available for ArsB, precluding the molecular understanding of substrate recognition and electrochemical coupling in the metalloid transporter.

Structures of other IT proteins reveal a conserved architecture composed of a scaffold domain and a transport domain^23,24^. The latter binds and transports the substrate by moving along the membrane relative to the scaffold domain via an ‘elevator-type mechanism’, alternating between inward and outward conformations^12,25^. This motion of the transport domain is correlated with the movement of the substrate-binding site itself, thus providing ‘alternating-access’ to the substrate between the two sides of the membrane. The elevator-type mechanism is well defined for secondary symporters of the IT superfamily, as guided by Na+/substrate-bound structures^12,13,26^. In contrast, how secondary antiport operates in the framework of the elevator-type mechanism remains elusive. In this light, ArsB can serve as a model for understanding antiport mechanisms within the broader IT superfamily.

In this work, we structurally characterized ArsB using single particle electron cryo-microscopy (cryo-EM) from the intrinsically metalloid-tolerant species, *Leptospirillum ferriphilum* strain ML-04. The structures reveal an ‘inward-facing’ conformation of ArsB in apo and As^III^/Sb^III^-bound states. Supported by mutagenesis and growth assays in the presence of As^III^, the structures establish the molecular basis for metalloid recognition and transport. The metalloid interacts with polar residues in a conserved pocket via hydrogen-bonding interactions. Moreover, we demonstrate that As^III^ resistance conferred by ArsB to *E. coli* varies with external pH, consistent with H^+^-gradient dependence of metalloid transport. Asp112 and Asp148 near the metalloid-binding pocket may act as protonation sites, enabling proton shuttling across the membrane to drive ‘H^+^/metalloid antiport’. Overall, this work uncovers the structural basis of metalloid recognition and H^+^-coupling in ArsB; these findings are key steps toward a comprehensive understanding of the metalloid efflux mechanism of the transporter.

## Results

### Cryo-EM structure of *Lf*ArsB in apo state

We identified an ArsB ortholog from the thermotolerant and acidophilic bacterium, *L. ferriphilum* strain ML-04 (*Lf*ArsB)^27^, and recently structurally characterized the corresponding ArsA ATPase^28^. *Lf*ArsB shows 75% sequence identity to *E. coli* ArsB (*Ec*ArsB), for which functional assays have been previously performed^8^. As *L. ferriphilum* is recognized for its high arsenic tolerance^27^, we performed structural characterization of *Lf*ArsB in this work to understand its metalloid transport mechanism.

Challenges with stable overexpression of ArsB have precluded its biochemical and structural characterization so far^29^. We accomplished the overexpression of *Lf*ArsB in *E. coli* using a C-terminal fusion to superfolder green fluorescent protein (GFP) (Supplementary Fig. 1a), inspired by a strategy proposed by Hsieh and coworkers^30^. Since topology prediction of ArsB determined both N- and C-termini to localize to the periplasm^31^, a single-pass transmembrane domain, glycophorin A (GpA) was inserted between ArsB and GFP to ensure that GFP localized to the cytoplasm. Additionally, an HRV-3C protease cleavage site flanked by Gly-Ser (GS) linkers was inserted between ArsB and GpA sequences to allow removal of the tag during purification of ArsB. Finally, an 8x-His tag, separated by a 10x-GS linker, was attached after GFP to enable affinity purification. The fusion construct did not interfere with the function of ArsB, as demonstrated by comparable levels of As^III^ resistance conferred to *E. coli* AW3110 (Δ*ars*) cells by both wild-type and the corresponding GFP-fusion construct of *Ec*ArsB (Supplementary Fig. 1b). The GFP-fusion construct enhanced the yield of ArsB and allowed for tracking the protein during purification. ArsB was purified using Ni^2+^-affinity chromatography and size exclusion chromatography (SEC). Two distinct peaks, designated P1 and P2, were observed on the SEC trace when ArsB was purified in n-dodecyl-β-D-maltoside (DDM) (Supplementary Fig. 2a). Preliminary cryo-EM analysis of both P1 and P2 samples revealed transmembrane domains (TMDs) enclosed in detergent micelles that appeared to correspond to dimeric and monomeric species, respectively (Supplementary Fig. 2b, c).

We obtained a high-resolution cryo-EM reconstruction of the ArsB dimer from the P1 sample of ArsB solubilized in lauryl maltose neopentyl glycol/cholesteryl hemisuccinate (LMNG/CHS) (Supplementary Figs. 3a-c). The size of the ArsB monomer (45 kDa) is borderline for characterization by conventional cryo-EM methods, while the dimer is of sufficient size to obtain high resolution. Moreover, segments of ArsB TMDs protruding out of the detergent micelles apparent in the 2D classes (Supplementary Fig. 3c) likely facilitated particle alignment to enable high-resolution reconstruction. From a cumulative dataset of 11,686 movies collected on a 300 keV Titan Krios TEM at a nominal magnification of 130,000x, two populations of ArsB dimer with distinct two-fold symmetry axes were reconstructed (Supplementary Figs. 4 and 5a, b; Supplementary Table 1). An AlphaFold3 model of the *Lf*ArsB monomer was docked into the maps, and the final models were built through iterative refinement (Supplementary Fig. 6). Of the two dimer states, one is a ‘parallel dimer’ resolved to 3.6 Å overall resolution with an apparent two-fold rotation axis perpendicular to the expected plane of the membrane, and the monomers are oriented so that the respective termini are localized to the same side of the micelle (Supplementary Fig. 7a). The second is an ‘antiparallel dimer’ resolved to 3.1 Å overall resolution with an apparent two-fold axis within the expected membrane plane, and the respective termini of each monomer localized to opposite sides of the micelle (Supplementary Fig. 7b).

The individual subunits superimpose closely between the two dimeric states (C_⍺_ root mean square deviation (r.m.s.d.) = 0.37 Å) allowing a simple description of an ArsB monomer (Fig. 1a). The monomer is composed of a total of 10 TMDs and 2 sets of hairpin helices with both N- and C-termini localized to the periplasmic side (Fig. 1b). ArsB adopts an inverted repeat fold composed of the N-terminal half (TM1-5) and the C-terminal half (TM6-10), where the two halves are structurally homologous (Fig. 1b). The two halves are connected by a linker sequence (Arg199-Val221) on the cytoplasmic side. A similar topology is found in other transporters of the IT superfamily^15,18,26^. The TMDs are organized into a scaffold domain and a transport domain, an architecture typical of elevator-type transporters (Fig. 1c). A helix-turn-helix hairpin motif (HP1) is formed by helices HP1a and HP1b and the discontinuous helix 4 is formed by TM4a and TM4b, with a break at loop L1 in the N-terminal half (Fig. 1d). Likewise, in the C-terminal half, a hairpin motif (HP2) is formed by helices HP2a and HP2b and the discontinuous helix 9 is formed by TM9a and TM9b, with a break at loop L2 (Fig. 1d). These two motifs, hereafter denoted as ‘hairpin-loop motifs’, form the core of the transport domain. Moreover, the scaffold domain is composed of TM1-3 of the N-terminal half and TM6-8 of the C-terminal half, where helical segments 3c and 8c are oriented horizontally or parallel to the expected membrane plane on the opposite ends of ArsB (Fig. 1c, d). The interface of the scaffold and transport domains is largely composed of hydrophobic residues (Supplementary Fig. 8), as observed in other IT proteins^13,15,16^. Interactions at this interface support low-barrier ‘elevator-type’ conformational changes that may facilitate metalloid transport.

**Figure 1.**
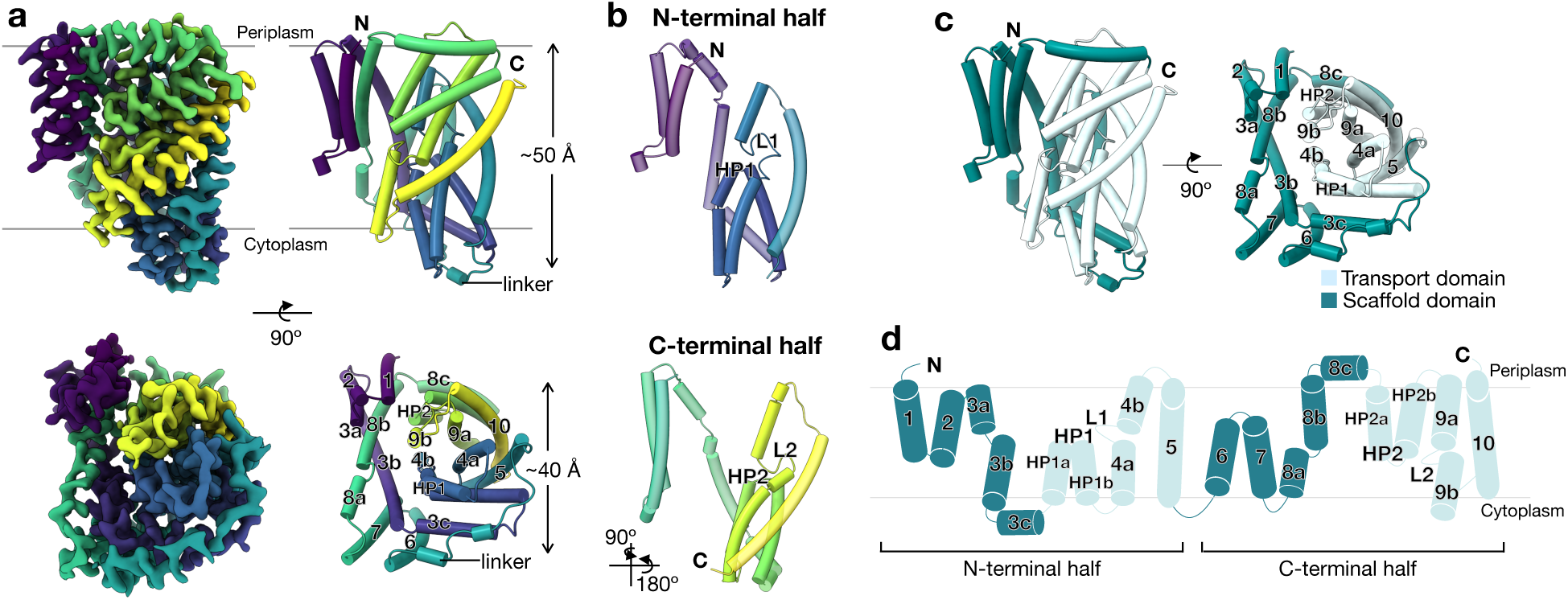
Cryo-EM structure of *Lf*ArsB in apo state. **a** Sharpened cryo-EM map (3.1 Å) (left) and cartoon representation of the model (right) as viewed from the membrane plane (top) and the cytoplasmic side (bottom). Both map and model are colored by transmembrane domains (TMDs) using a viridis coloring palette^50^. **b** Cartoon representation of N- and C-halves highlighting the conserved topology and the hairpin-loop (HP1/2) and loop motifs (L1/2) in the two halves. **c** Two perpendicular orientations of *Lf*ArsB model highlighting the scaffold (teal) and transport domain (cyan) organization. **d** Membrane topology diagram showing the arrangement and numbering of the TMDs.

### Metalloid-binding site of ArsB

To characterize the metalloid-binding site, we determined the cryo-EM structure of ArsB in the presence of 2 mM As^III^ (Supplementary Figs. 9 and 5c; Supplementary Table 1). Both parallel and antiparallel dimer reconstructions were obtained, as seen in the apo ArsB dataset. Additionally, for this dataset, 3D classification resolved a tetrameric complex, a dimer of dimers of ArsB, consisting of both the parallel and antiparallel dimers (Supplementary Fig. 9). Although higher-order oligomers were also indicated in the 2D class averages of the apo dataset, these could not be resolved in the 3D reconstruction. We believe that the variable oligomeric states in these structures are artefacts of detergent purification. Within the tetramer, the parallel dimer yielded the highest-quality reconstruction. To observe high-resolution features within the parallel dimer, we masked the poorly resolved inverted subunits of the antiparallel dimer from the map and performed focused refinement on the parallel dimer, yielding a final reconstruction at 3.2 Å overall resolution (Supplementary Fig. 5c). The model was built and refined into this map. Overall, the resulting monomer structure was very similar to the apo ArsB structure (C_⍺_ r.m.s.d. = 0.39 Å).

The metalloid-binding site was identified in a small pocket at the core of the transport domain enclosed by the HP1-L1 motif of the N-half, and the HP2-L2 motif of the C-half (Fig. 2a). Closer inspection of this pocket in the cryo-EM map revealed additional density that was not observed in the apo map (Supplementary Fig. 10). We modeled this density with an As(OH)_3_ moiety (pK_a1_ = 9.2), which is the thermodynamically-dominant As^III^ species at physiologically-relevant pH (Fig. 2b)^22^. The As^III^ -binding site is composed of polar residues located at the helix turns of HP1 and HP2, and L1 and L2 loops, forming hydrogen-bonding contacts with As(OH)_3_. The sidechains of Asn111 on HP1, Asn337 on HP2 and Ser380 on L2 are positioned within 3 Å of the hydroxyl groups (Fig. 2b). The sidechains of Asp112 on HP1 and Asn158 on L1, and the backbone amino group of Ala382 on L2 are also located within polar interaction distance (<4 Å) from As(OH)_3_ (Fig. 2b). The arrangement of non-ionizable polar residues in the binding pocket is consistent with the binding of a trigonal As(OH)_3_ species bearing no overall charge; this setup dictates the substrate specificity of ArsB. Moreover, the polar pocket is surrounded by hydrophobic residues including Phe60 from TM3b of the scaffold domain and Leu159, Val160, Leu381, and Met338 of the transport domain (Fig. 2b). These residues form van der Waals contacts with As(OH)_3_ and may collectively strengthen the polar interactions between the metalloid and the surrounding residues by forming a low dielectric environment in the metalloid-binding pocket. Intriguingly, the density for Phe60 sidechain appears to be disordered and shifted toward As(OH)_3_ in the metalloid-bound structure, in contrast to the apo structure, suggesting multiple conformations of the sidechain (Supplementary Fig. 11). In DASS proteins, this conserved phenylalanine flips between two rotamers, functioning as a gating mechanism that coordinates substrate binding and transport^20,32^.

**Figure 2.**
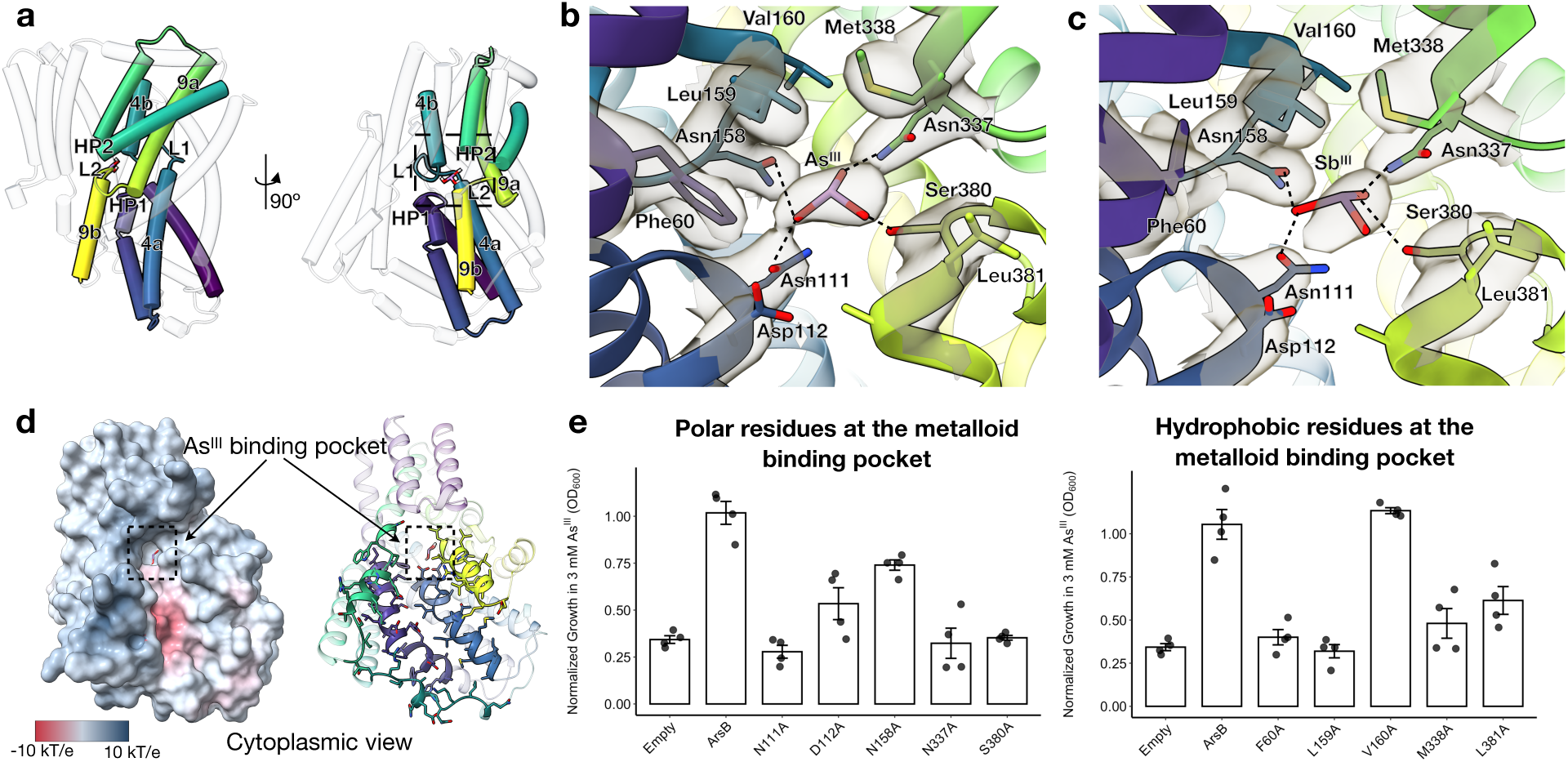
Metalloid recognition in *Lf*ArsB. **a** Cartoon representation in two orientations showing the position of the metalloid-binding pocket enclosed by TMDs of the transport domain. **b** Interactions between arsenite (As(OH)_3_) and residues in the metalloid-binding pocket. **c** Interactions between antimonite (Sb(OH)_3_) and residues in the metalloid-binding pocket. For both **b** and **c**, Coulomb potential map density is shown for the metalloid and neighboring residues. Hydrogen bonds between the metalloid and neighboring residues are shown with dotted lines connecting donor-acceptor atoms. H atoms of the metalloid hydroxyl groups are not shown. **d** Cytoplasmic view of As^III^-bound ArsB structure as electrostatic potential surface (left) and cartoon (right) representations. The As^III^-binding pocket is located at the end of a funnel-shaped cavity exposed to the cytoplasm **that** is lined by polar residues with negative electrostatic potential (red surface). Adaptive Poisson-Boltzmann Solver (APBS) was used to calculate the electrostatic potential surface^51^. **e** Growth of *E. coli* AW3110 cells (Δ*ars*) complemented with wild-type *Ec*ArsB or point mutants of the polar residues (left) and hydrophobic residues (right) at the metalloid-binding pocket in the presence of 3 mM As^III^. OD_600_ values after 10-h growth in the presence of As^III^ are reported after normalizing by OD_600_ values in absence of As^III^. Biological quadruplicates are reported for each sample and error bars represent standard error of mean.

To validate metalloid binding at this site, we determined the cryo-EM structure of ArsB in the presence of 2 mM Sb^III^ at 3.1 Å overall resolution (Supplementary Figs. 12 and 5d; Supplementary Table 1), and observed density corresponding to Sb(OH)_3_ at the same site with sidechain interactions preserved (Fig. 2c). Examination of the electrostatic surface of ArsB bound to As^III^ reveals that the binding pocket is accessible from the cytoplasmic side via a funnel-shaped cavity. The cavity leading to the metalloid-binding site, shown as an electronegative potential surface, is lined with polar residues (Fig. 2d). These observations indicate that the state of ArsB seen in these structures is accessible to the metalloid substrate from the cytoplasm.

To support the interactions observed at the metalloid-binding site in our cryo-EM structures, we performed alanine mutagenesis of the binding-site residues of ArsB and measured the growth of AW3110 cells complemented with ArsB variants under toxic As^III^ concentrations. These assays were performed using *Ec*ArsB, where the metalloid-binding pocket residues are conserved relative to *Lf*ArsB (Supplementary Fig. 13). In the presence of 3 mM As^III^, cells bearing wild-type *Ec*ArsB were fully resistant to As^III^, while those bearing the alanine mutants of Asn111, Asn337, and Ser380 showed significantly reduced As^III^ tolerance. This finding is consistent with the As^III^-bound structure, which shows direct contacts between these sidechains and As^III^ (Fig. 2e, left). D112A and N158A show intermediate resistance, consistent with marginal interactions between these sidechains and As^III^ (Fig. 2e, left). Additionally, mutations in hydrophobic residues surrounding the As^III^-binding site largely resulted in a significant decrease in resistance (Fig. 2e, right). Altogether, these functional assays corroborate the interactions inferred at the metalloid-binding site in the ArsB structure, which either stabilize metalloid binding or facilitate subsequent metalloid translocation.

### Structural comparison of ArsB with DASS family transporters

Comparison of the ArsB cryo-EM structures to homologs of the DASS family revealed a conserved architecture of the transmembrane helices characteristic of the IT superfamily fold (Fig. 3a). This ArsB conformation resembles the ‘inward-open’ conformation observed in the structures of the Na^+^-dependent dicarboxylate transporter from *V. cholerae* (*Vc*INDY) and the Na^+^-dependent citrate transporter from humans (*Hs*NaCT) (Fig. 3a, left). In each of these structures, the transport domain helices extend out of the membrane defined by the horizontal helices of the scaffold domain (helices 3c and 8c in ArsB; Fig. 1a) on the cytoplasmic side, whereas the scaffold domain is confined within the membrane. A notable difference between the ArsB and DASS protein structures is the position of TM9b in the transport domain relative to the scaffold domain. In the ArsB structure, TM9b, along with the upstream L1 motif, appears to shift closer to the scaffold domain and the substrate-binding pocket compared to the analogous motifs in *Vc*INDY and *Hs*NaCT structures (Fig. 3a, right). The position of L1 and TM9b motifs in ArsB more closely resembles that of the ‘inward-occluded’ conformation of INDY from *Drosophila melanogaster* (*Dm*INDY) bound to its substrate citrate (Fig. 3b)^32^. Consequently, the substrate-binding pocket is more compact in the ArsB and *Dm*INDY structures (Supplementary Fig. 14a). The respective substrates of ArsB and DASS proteins bind at similar binding pockets stabilized by polar interactions with residues of the hairpin-loop (HP1-L1 and HP2-L2) motifs of the transport domain (Fig. 3c, d and Supplementary Fig. 14b). The difference in the size of the substrate-binding pocket between ArsB and NaCT corelates with a smaller spacing in the hairpin-loop motifs, where a difference of ∼2 Å shrinks the binding pocket in ArsB (Fig. 3c).

**Figure 3.**
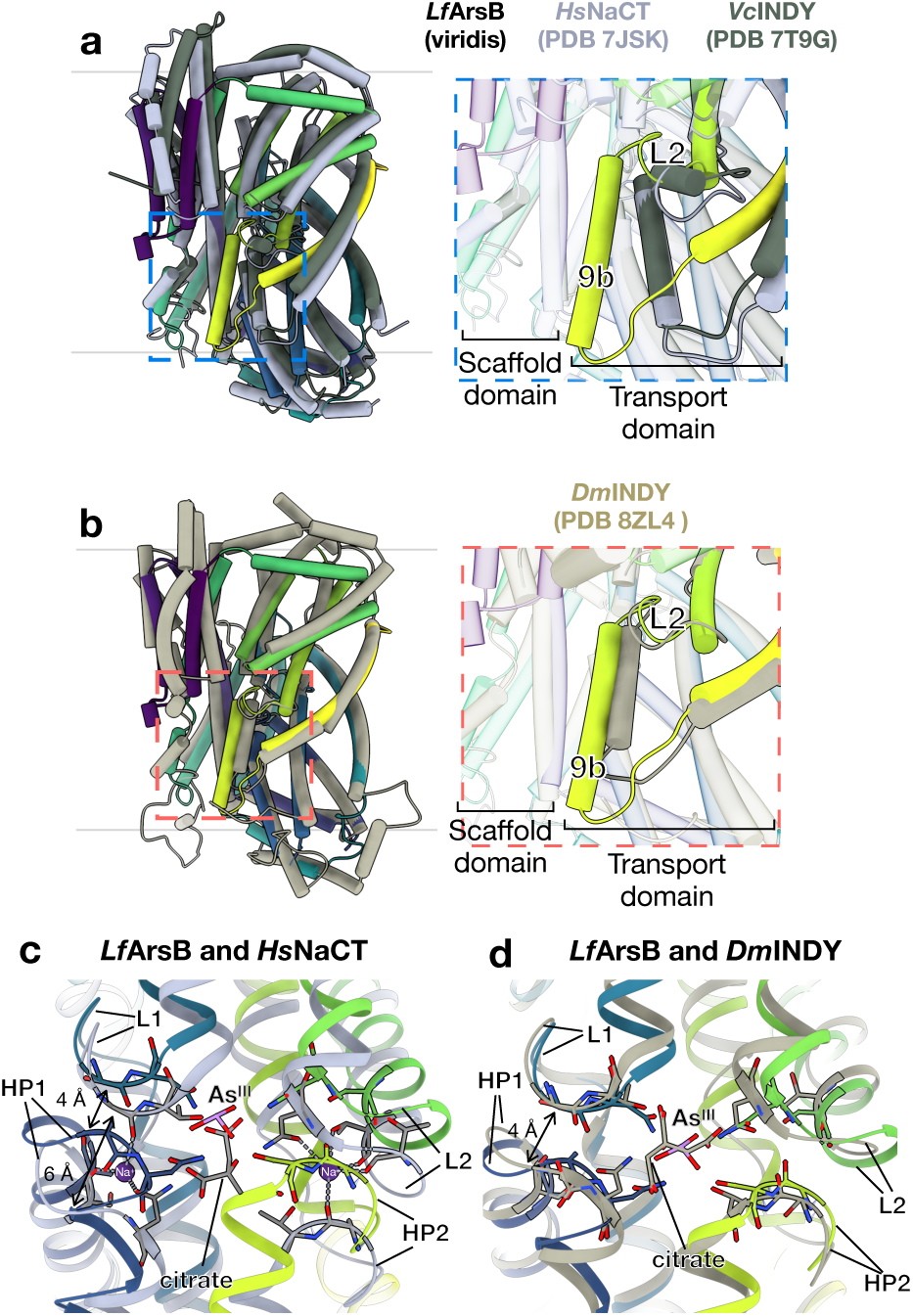
Structural comparison of *Lf*ArsB with DASS family transporters. **a** As^III^-bound *Lf*ArsB structure (viridis) superposed on the ‘inward-open’ state of *Hs*NaCT (gray) and *Vc*INDY (dark green). On right, zoomed-in view of structural changes in the transport domain relative to the scaffold domain between ArsB and the two DASS transporters. **b** As^III^-bound *Lf*ArsB structure (viridis) superposed on the ‘inward-occluded’ state of *Dm*INDY (light brown). On right, zoomed-in view of the structural changes in the transport domain relative to the scaffold domain between ArsB and *Dm*INDY. **c, d** Comparison of the substrate-binding site and the hairpin-loop motifs (HP1-L1 and HP2-L2) of As^III^-bound *Lf*ArsB structure with citrate/Na^+^-bound *Hs*NaCT ‘inward-open’ structure (**c**), and with citrate-bound *Dm*INDY ‘inward-occluded’ structure (**d**).

A key difference between ArsB and DASS transporters lies in their coupling-ion specificity. DASS proteins typically utilize Na^+^-coupling for substrate transport^13,14^, whereas ArsB exploits the H^+^-gradient across the membrane^7,8^. Na^+^ ions intercalate in the cavity formed by the HP1-L1 and HP2-L2 motifs in DASS proteins (Fig. 3c), stabilized by backbone carbonyls and polar sidechains of residues in these motifs. However, despite the conservation of these Na^+^-coordinating residues and the presence of 100 mM NaCl in the sample buffer (Supplementary Fig. 14b), no density corresponding to a monovalent cation was observed in the cavity formed by the hairpin-loop motifs in either the apo or the metalloid-bound ArsB structures. A more constricted cavity formed at the HP1-L1 and HP2-L2 motifs in ArsB, compared to DASS transporters such as *Hs*NaCT, may no longer support Na^+^ binding (Supplementary Fig. 14b). This phenomenon is also observed in the ‘inward-occluded’ state of *Dm*INDY, defined by a smaller substrate-binding pocket (Fig. 3d)^32^. As both ArsB and *Dm*INDY use H^+^ instead of Na^+^ as coupling ion, this structural feature is consistent with a shift in coupling ion preference to H^+^, thereby indicating an alternative mechanism of ion coupling. Since a fully ‘inward-open’ conformation of ArsB similar to that of *Vc*INDY and *Hs*NaCT is not adopted in our structures, even in the apo state, the conformation observed here is defined as the ‘inward-facing’ state that enables metalloid binding from the cytoplasmic side before undergoing elevator-type movements for metalloid translocation.

### H^+^-coupling mechanism of ArsB

While Na^+^ binding and coupling are well-characterized in DASS transporters, the structural basis and mechanism for H^+^-coupled metalloid efflux in ArsB remain unclear. To understand the relationship between the H^+^-gradient and metalloid efflux, we measured As^III^ resistance in AW3110 cells complemented with *Ec*ArsB at different external pH levels. Cells with *Ec*ArsB were grown at various As^III^ concentrations up to 10 mM across pH values from 5.0 to 8.0 in 1-unit increments, adjusted by buffering the LB media. Although cells tolerated As^III^ at all pH values, higher resistance was observed at lower (more acidic) external pH values (Fig. 4a). This pH-dependent growth in the presence of As^III^ was not seen in cells with an empty vector (Supplementary Fig. 15), confirming that the effect is due to ArsB-mediated As^III^ efflux. Altering the external pH affects the ΔpH across the membrane, considering that *E. coli* maintains a stable intracellular pH between 7.2 and 7.8^33,34^. Since ΔpH is positive at moderately acidic external pH, this result supports the H^+^-gradient dependence of ArsB.

**Figure 4.**
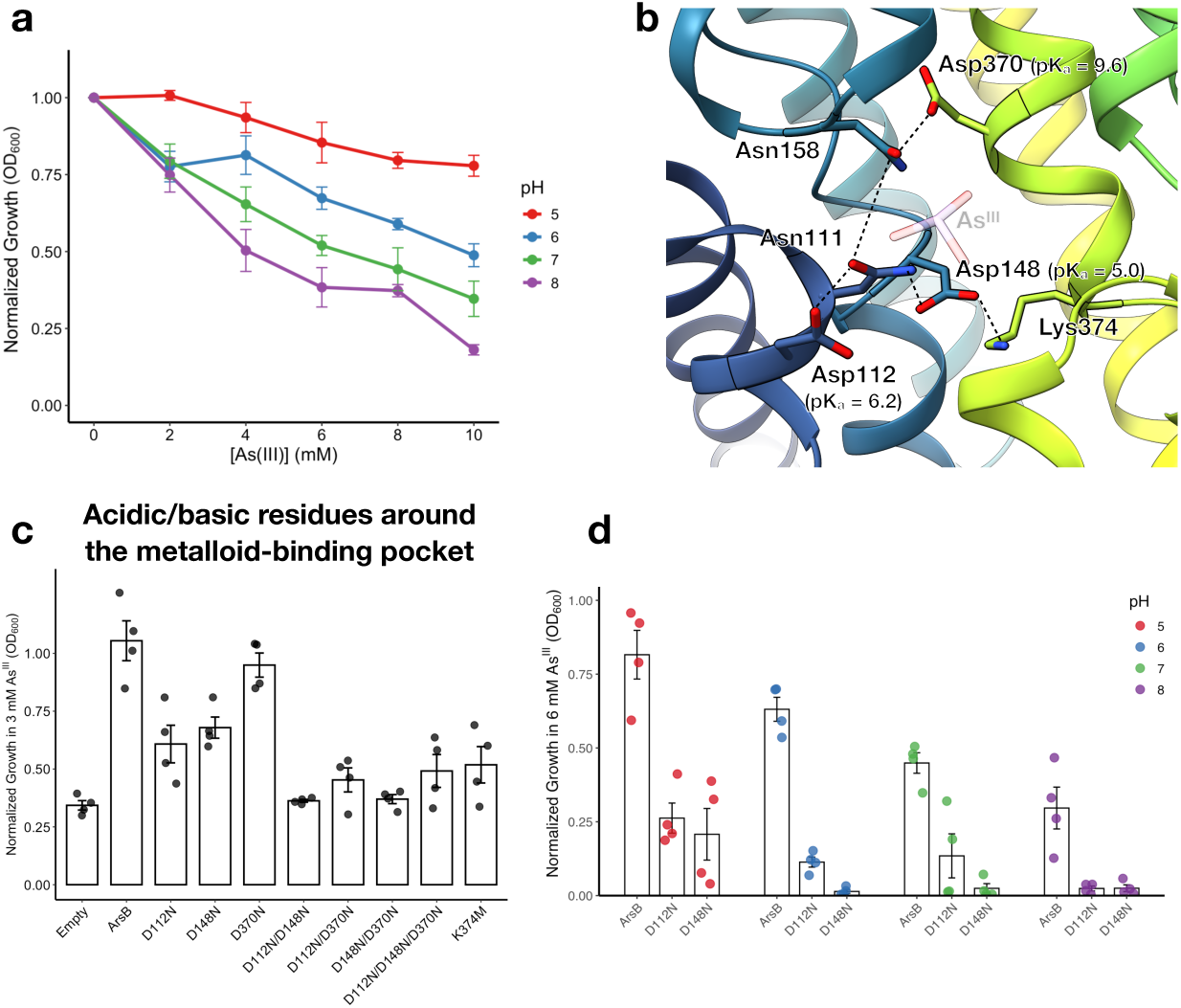
H^+^-coupled metalloid transport by ArsB. **a** External pH-dependence of As^III^ resistance conferred by *Ec*ArsB in AW3110 cells. **b** Putative H^+^-coupling residues (Asp112, Asp148 and Asp370) at the metalloid-binding pocket of *Lf*ArsB apo structure shown as sticks. Corresponding predicted pK_a_ values calculated using PROPKA3^37^ are shown in parentheses. Neighboring residues forming polar interactions with these aspartates are shown as sticks as well. Position of the metalloid substrate (As(OH)_3_) at the binding site is shown at 50% opacity for reference. **c** Normalized growth in presence of 3 mM As^III^ for *Ec*ArsB variants of the putative H^+^-coupling residues. **d** External pH-dependence of cell growth with wild-type *Ec*ArsB, D112N and D148N in presence of 6 mM As^III^. For growth data in panels **a**, **c** and **d**, biological quadruplicates are reported for each sample, and the error bars represent standard error of mean.

We aimed to understand the molecular basis of H^+^-coupling in ArsB through our cryo-EM structures. H^+^-coupling in other secondary transporters, such as the *S. aureus* multidrug efflux pump NorA and the mitochondrial calcium transporter NCLX, has been shown to involve ionizable acid/base residues (Asp/Glu)^35,36^. Examining the metalloid-binding pocket of ArsB, we identified three conserved ionizable residues—Asp112 (HP1), Asp148 (TM4a), and Asp370 (TM9a) (Fig. 4b)—as potential H^+^-coupling sites that may undergo protonation and deprotonation to facilitate H^+^/metalloid antiport. Among these, Asp148 forms a salt bridge with Lys374 on TM9a. Notably, these residues are not found in Na^+^-dependent DASS transporters (Supplementary Fig. 16a). To assess their functional roles, we performed As^III^ resistance growth assays on *Ec*ArsB variants with these aspartates replaced by asparagines. The rationale for the asparagine mutation was to disrupt the ionizable nature of the carboxylate side chain while maintaining polar interactions with surrounding residues, thereby ensuring that the observed effects of the mutation were likely due to the disruption of the H^+^-coupling site. Mutations of Asp112 and Asp148 impaired growth in 3 mM As^III^, while mutation of Asp370 had only a marginal effect (Fig. 4c). Additionally, mutating Lys374 to methionine disrupted growth, likely due to breaking the salt bridge with Asp148 (Fig. 4c). Double and triple aspartate mutants almost completely abolished growth, supporting a collective role for these residues (Fig. 4c). In our ‘inward-facing’ ArsB structures, Asp370 is buried within the hydrophobic core of the TMDs, while Asp112 is located near the end of the cavity that exposes the metalloid-binding site to the cytoplasm, and Asp148 is on the cytoplasmic side next to Lys374 (Fig. 4b and Supplementary Fig. 16b). Assuming the solvent accessibilities of these aspartates remain relatively unchanged during transport, as expected for elevator-type transporters where the transport domain undergoes rigid-body movements along the membrane^25^, we inferred that Asp112 and Asp148 could serve as H^+^-coupling residues. This hypothesis is supported by pK_a_ predictions of the carboxylate sidechains of Asp112 (6.2), Asp148 (5.0), and Asp370 (9.6) in our apo ArsB structure using the empirical pK_a_ calculation package PROPKA3 (Fig. 4b)^37^. In the ‘inward-facing’ state observed, the metalloid-binding pocket faces the cytoplasm (pH ∼7.2-7.8). Under these conditions, Asp370 is likely protonated, while Asp112 and Asp148 are deprotonated, facilitating H^+^ transport into the cytoplasm. The role of Asp112 and Asp148 is consistent with the growth assays for their asparagine mutants (Fig. 4c). Further, we compared the growth of cells expressing D112N and D148N variants to wild-type ArsB at different external pH levels under 6 mM As^III^. Consistent with their proposed roles in H^+^-coupling, both mutants showed poor growth across all tested pH values (Fig. 4d).

## Discussion

As a metalloid-specific efflux pump, ArsB plays a central role in bacterial survival in arsenic-contaminated environments. In this work, we solved high-resolution cryo-EM structures of *Lf*ArsB in both apo and metalloid-bound states. In the framework of the IT superfamily fold, these structures represent an ‘inward-facing’ conformation that is competent for metalloid capture from the cytoplasm. Metalloid recognition is primarily enabled by hydrogen-bonding interactions with polar residues located on the HP1-L1 and HP2-L2 motifs in the metalloid-binding pocket. This binding mode resembles the divalent-substrate binding in DASS transporters (Supplementary Fig. 14b), yet is compatible with binding to neutral As(OH)_3_ or Sb(OH)_3_ species. Similarly, in GlpF, which facilitates passive As(OH)_3_ import in bacteria, hydrogen-bonding contacts with asparagine and backbone atoms enable substrate recognition^38^. This binding mode contrasts with the three-coordinate As-S linkage formed between As^III^ and cysteine-rich proteins, including ArsA and the As^III^-metallochaperone ArsD, that require stronger polar covalent interactions to mediate metalloid capture from the cytoplasm^28,39^. Notably, for both ArsB and GlpF, low-energy interactions with neutral metalloids are likely advantageous for effective translocation across the membrane.

ArsB is a H^+^-coupled antiporter that exploits the inward-directed H^+^-gradient to drive outward-directed metalloid transport, in contrast to many other IT proteins that perform Na^+^-coupled substrate symport^14,18,23^. Unlike the anionic substrates of these transporters, a neutral metalloid substrate does not require Na^+^-mediated charge compensation to facilitate transport. Consistent with this, our structural analysis supports the preference for H^+^ as the coupling ion in ArsB. A more compact substrate-binding pocket, observed in our ‘inward-facing’ structure compared with DASS transporters such as NaCT (Fig. 3c), makes Na^+^-binding between the hairpin-loop motifs less likely. Additionally, we identified two functionally important residues – Asp112 and Asp148 – that are located at sites distinct from the hairpin-loop cavity where Na^+^ typically binds in DASS proteins. These ionizable residues may serve as potential H^+^-coupling sites to facilitate H^+^ influx during the transport cycle.

H^+^/substrate antiport mediated by ArsB is a novel mechanistic theme among elevator-type transporters of the IT superfamily. The ‘inward-facing’ conformation in our structures represents the initial stage of the ArsB transport cycle. This state may bind metalloids from the cytoplasm and, concomitantly, enable deprotonation of the conserved H^+^-coupling aspartates. The metalloid-bound transporter then may undergo elevator-type movements, transitioning into the ‘outward-facing’ conformation compatible with metalloid release into the periplasm. Protonation of the H^+^-coupling aspartates in the ‘outward-facing’ state may facilitate H^+^ influx as the transporter transitions to the ‘inward-facing’ state, and hence, resets ArsB for another transport cycle. Structural insights into an ‘outward-facing’ state will further support this model for the H^+^/metalloid antiport mechanism.

The structures of ArsB reported here reveal both parallel and antiparallel dimeric architectures. Fortuitously, the two distinct dimeric states enabled us to solve the structure of the 45 kDa ArsB monomer without fiducial markers. Given the flipped orientation of monomers between these two dimers and the relatively sparse interface in the parallel dimer (Supplementary Fig. 17a), we believe these oligomers are not physiological and are likely formed upon reconstitution of ArsB from the lipid membrane into detergent micelles. We note that formation of two such distinct dimers has also been observed for TRAP transporters in detergent micelles (Supplementary Fig. 17b)^19^. DASS transporter structures are dimeric, whereas both dimers and monomers have been reported for TRAP transporters^13,19^; however, the subunits within these dimers are believed to function independently^40^. In light of these observations, the functional unit of ArsB in the membrane is likely a monomer; nonetheless, the biological oligomeric state of ArsB remains to be experimentally determined.

Beyond H^+^-coupled secondary transport, certain ArsB orthologs, including those from *E. coli* and *L. ferriphilum*, can couple metalloid transport to ATP hydrolysis by associating with ArsA ATPase, thereby enhancing efflux efficiency. We previously showed that AlphaFold3 predictions for ArsA and ArsB, even in the presence of nucleotides and metalloid, do not support the formation of a stable complex^28^. However, in every predicted state, the metalloid-binding site of ArsA docks near the cytoplasmic side of ArsB, which is consistent with metalloid transfer. Based on this, we suggest that a low-affinity or transient interaction between the two proteins is possible. While a transient interaction might be beneficial for the vectorial transfer of metalloid from ArsA to ArsB, the structural mechanism behind this requires experimental validation. Due to its apparent mechanistic versatility, ArsB serves as a model system for understanding toxic metalloid efflux. This work offers a structural foundation for further mechanistic studies of ArsB-mediated metalloid transport, which would be valuable for the development of arsenic bioremediation strategies^9,10^.

## Methods

### Cloning of *Lf*ArsB construct

*The Lf*ArsB gene was obtained from Twist Bioscience (South San Francisco, CA) and cloned into the multiple-cloning site 2 (MCS-2) of the pETDuet-1 backbone. DNA amplification and cloning were performed using Q5^®^ High-Fidelity 2x Master Mix and NEBuilder^®^ HiFi-DNA Assembly Master Mix, respectively. Expression of ArsB was successfully achieved using a GFP-fusion-based strategy reported by Hsieh and coworkers^30^. To construct the ArsB-GFP fusion, GpA-GFP-8xHis was added to the C-terminus of *Lf*ArsB. A Human Rhinovirus (HRV)-3C protease cleavage site flanked by an upstream 7xGS linker and a downstream 2xGS linker was inserted between ArsB and GpA sequences to engineer a cleavable construct. Additionally, a 10xGS linker was inserted between GFP and 8xHis sequences. The final expression construct constituted *Lf*ArsB-7xGS-HRV-3C-2xGS-GpA-GFP-10xGS-8xHis in the MCS-2 of pETDuet-1 backbone.

### Expression and purification of *Lf*ArsB

*The Lf*ArsB expression plasmid was transformed into *E. coli* Nico21(DE3) competent cells (NEB). Cells from lysogeny-broth (LB) agar plate grown overnight were inoculated into a starter culture containing 100 µg/mL ampicillin and grown as a starter culture at 37 °C for 2 hours before inoculating into 2x yeast-tryptone (2xYT) media in 1-L flasks. Cells were induced with 0.4 mM isopropyl-β-D-thiogalactopyranoside (IPTG) at an optical density of 0.6-0.8 and grown at 18 °C for 16-20 hours. Successful expression was indicated by a pale-yellowish appearance of the pellet owing to the ArsB-GFP fusion. Cells were harvested and lysed by passing 3x times through an M-110L microfluidizer (Microfluidics). The lysate was clarified by centrifugation at 24,000 xg at 4 °C for 30 min, followed by isolation of the membrane fraction by centrifugation at 38,000 rpm at 4 °C for 40 min using an Optima XE-90 Ultracentrifuge (Beckman Coulter). The resulting membrane pellet was resuspended in extraction buffer consisting of 50 mM 4-(2-hydroxyethyl)-1-piperazineethanesulfonic acid (HEPES) at pH 7.5, 300 mM NaCl, 5% glycerol, 1%/0.1% LMNG/CHS, 10 mM imidazole, and 5 mM β-mercaptoethanol, and incubated at 4 °C for 1 hour. The supernatant was incubated with 1 mL Ni-nitrilotriacetic acid (Ni-NTA) resin (Qiagen) pre-equilibrated with extraction buffer at 4 °C for 1 hour. Protein-bound resin, with a bright green appearance due to GFP, was spun down at 700 xg at 4 °C for 5 min, and then washed with 50 column volumes (CV) of wash buffer (50 mM HEPES pH 7.5, 300 mM NaCl, 5% glycerol, 0.005%/0.0005% LMNG/CHS, 30 mM imidazole and 5 mM β-mercaptoethanol) and one CV fractions were collected with elution buffer (wash buffer + 200 mM imidazole). Purity of the fractions was assessed by sodium dodecyl sulphate-polyacrylamide gel electrophoresis (SDS-PAGE) analysis. The protein eluted in the first four fractions with high purity. These fractions were pooled and concentrated to 3 mL using a 100 kDa molecular weight cut-off (MWCO) Amicon filter (Milli-pore) and then exchanged into low-salt and imidazole-free buffer or the SEC buffer (50 mM HEPES pH 7.5, 100 mM NaCl, 5% glycerol, 0.005%/0.0005% LMNG/CHS, and 5 mM β-mercaptoethanol) using a 10-mL desalting column (Bio-Rad), in preparation for protease cleavage. Purified HRV-3C protease was added at a ratio of 4 mg per mL of Ni eluant and incubated at 4 °C for 16 h. Fully cleaved ArsB was separated from cleaved GpA-GFP-His fusion tag, uncleaved ArsB, and His-tagged 3C protease by passing again through the Ni-NTA resin. The total cleaved ArsB sample was mixed with Ni-NTA resin pre-equilibrated with SEC buffer, flow-through was collected, and the process was repeated for a total of four passes. Two additional fractions were collected using SEC buffer supplemented with 30 mM imidazole. After the purity of the fractions was assessed by SDS-PAGE, fractions containing only cleaved ArsB were pooled and concentrated to 500 μL using a 100 kDa MWCO Amicon filter (Millipore). The sample was injected into a Superdex 200 Increase 10/300 GL column (Cytiva) pre-equilibrated with filtered and degassed SEC buffer, and the column was run at 0.4 mL/min, collecting 250 μL fractions. ArsB typically eluted as two distinct peaks around 11 mL (P1) and 12 mL (P2); a shoulder usually appeared before the 11-mL peak. The As^III^-bound structure was solved from ArsB extracted and purified in LMNG alone.

### Cryo-EM sample preparation

Fractions indicating the presence of pure and fully cleaved ArsB were pooled and concentrated to 4-6 mg/mL using Amicon 100-kDa concentration filter spun at 7500 xg at 4 °C. The molecular weight and extinction coefficient at 280 nm for cleaved ArsB were estimated as 47.2 kDa and 73,900 M^-1^ cm^-1^, respectively. For the apo structure, 3.0 μL sample, supplemented with 0.05% 3-[(3-cholamidopropyl)dimethylammonio]-2-hydroxy-1-propanesulfonate (CHAPSO), was applied to glow-discharged Quantifoil holey carbon R1.2/1.3 300 Mesh, Copper (Quantifoil, Micro Tools GmbH) grids using a Vitrobot (FEI Vitrobot Mark v4 x2, Mark v3). For metalloid-bound structures, the sample was mixed with 2 mM sodium arsenite or potassium antimonyl tartrate before CHAPSO was added. In each case, grids were blotted at 100% humidity and 4 °C followed immediately by plunge-freezing into liquid ethane.

### Cryo-EM data acquisition and processing

Cryo-EM data were collected on a 300 kV Titan Krios TEM equipped with a Gatan K3 direct electron detector and Gatan Energy Filter (slit-width of 20 eV) in super-resolution mode using SerialEM^41^. Datasets were acquired at 130,000x magnification with a raw pixel size of 0.325 Å pixel^-1^, electron exposure of 70 e^-^ Å^-2^ over 40 frames, and a defocus range of -0.8 to -2.8 μm in correlated double sampling (CDS) mode^42^. All datasets were processed in cryoSPARC v4.7.1^43^, using the same general processing workflow as described here, unless stated otherwise. Data collection and processing details for individual datasets are listed in Supplementary Table 1. During data acquisition, cryoSPARC LIVE was enabled for on-the-fly patch motion correction (0.5 F-cropping), contrast transfer function (CTF) estimation and manual curation of the collected movies. Motion-corrected micrographs having CTF outside the range of 2.5-5 Å were filtered out. Initial particles were picked using a Topaz model^44^ trained on a curated particle set from a dataset collected on DDM-solubilized ArsB sample. The particle set from DDM dataset was obtained by blob picking, extraction from micrographs and multiple rounds of 2D classification. Topaz-picked particles were extracted with a 2x bin (1.3 Å pixel^-1^) and subjected to one round of 2D classification to remove obvious junk particles or empty micelles. *Ab initio* reconstruction was performed on this particle set to obtain four 3D volumes. Volumes that showed TMD density were subjected to iterative *ab initio* reconstruction until no further junk particles could be removed. This curated set of particles was then used to train a new Topaz model, to re-extract particles from the micrographs, and perform iterative *ab initio* reconstructions. Once a high-resolution volume was obtained, particles were re-extracted with no binning (0.65 Å pixel^-1^) followed by a few more rounds of *ab initio* reconstructions. Reference-based motion correction, global or local CTF refinement and non-uniform refinement^45^ were performed to produce the final set of particles resulting in a high-resolution map. Finally, local refinement was performed using a soft mask for the ArsB dimer to obtain a high-quality final map. C2 symmetry was applied in each dataset. Overall map resolution was estimated using the gold-standard Fourier shell correlation (FSC) curve with a cutoff of 0.143 in cryoSPARC. For the ‘As^III^-bound parallel dimer’ dataset, the flipped orientation protomers in the tetramer were masked out and removed using ‘particle subtraction’ before the final round of local refinement was performed. Image processing pipelines for apo, As^III^-bound and Sb^III^-bound datasets are presented in Supplementary Figs. 4, 9 and 11 respectively.

### Model building and refinement

For each dataset, an initial model was built into the B-factor sharpened map from cryoSPARC using the AlphaFold3^46^ prediction of *Lf*ArsB as a reference. Backbone geometries and sidechain rotamers were refined using ‘Real-space refinement’ in Phenix v1.21.2^47^ and ‘Real-space refine zone’ in Coot v0.9.8.7^48^. For the As^III^ and Sb^III^-bound structures, geometry restraints for trigonal As(OH)_3_ and Sb(OH)_3_ species were generated using ‘elbow’ in Phenix. The ligand was docked in the density in Coot and refined using the restraints in Phenix.

### Arsenite resistance growth assays

The *Ec*ArsB gene with a C-terminal 6x-His tag was obtained from GenScript. *E. coli* strain AW3110 (Δ*ars*::cam), originally prepared by Carlin and coworkers^49^, was obtained as a gift from Prof. Chad Saltikov (University of California, Santa Cruz). In preparation for growth assays, AW3110 cells were first made electrocompetent. *Ec*ArsB-His and mutant genes were cloned into the MCS-2 of the pRSFDuet vector whose second T7 promoter upstream of the MCS-2 was deleted, resulting in the ‘pRSFdelT72-ArsB’ plasmid. Wild-type and mutant ArsB plasmids were transformed into AW3110 cells by electroporation. Transformed cells were plated onto LB agar plates containing 50 µg/mL kanamycin and 25 µg/mL chloramphenicol. Overnight cultures were grown from single colonies for ∼16 h. Growths were performed in 96-well plates in quadruplicates; 150 µL of LB media supplemented with 50 µg/mL of kanamycin and 25 µg/mL chloramphenicol, 0.1 mM IPTG, and increasing concentrations (mM) of sodium arsenite, were added to the wells.

Cells from the overnight cultures were added to the wells at 100-fold dilution. Growth was measured by recording the optical density at 600 nm (OD_600_) using a BioTek Epoch2 microplate reader at 37 °C, with cells shaking at 218 rpm (orbital mode). For the growth assays at different pH values, the LB medium was buffered with sodium citrate at pH 5.0, 3-(N-morpholino)propanesulfonic acid (MOPS) at pH 6.0 and 7.0, and HEPES at pH 8.0. OD_600_ values after 10 h of growth were recorded and normalized to the OD_600_ of samples with no arsenite. Normalized growth data were plotted in RStudio with assistance from ChatGPT.

## Data Availability

Atomic coordinates for all the structures reported in the paper are accessible on the Protein Data Bank (PDB) using the accession codes: 10TP (apo parallel dimer), 10TQ (apo antiparallel dimer), 10TU (As^III^-bound parallel dimer), and 10UA (Sb^III^-bound antiparallel dimer). Corresponding EM density maps can be accessed on the Electron Microscopy Data Bank (EMDB) using accession codes: EMD-75462 (apo parallel dimer), EMD-75463 (apo antiparallel dimer), EMD-75467 (As^III^-bound parallel dimer) and EMD-75468 (Sb^III^-bound antiparallel dimer).

## Acknowledgements

This work was supported by funding from HHMI (D.C.R.), Chan Zuckerberg Initiative (W.M.C.), and the Center for Environmental-Microbial Interactions at Caltech (S.M.). We acknowledge the Caltech Cryo-EM Resource Center supported by the Beckman Institute and are grateful to Dr. Songye Chen and Tyler Brittain for assistance with cryo-EM data collection. We thank Yen-Ju Lu for assistance with ArsB expression construct optimization. We are also grateful to Prof. Dianne Newman for her valuable feedback on the growth assays and providing access to the BioTek plate reader in the Newman lab. Finally, we thank Dr. Jacob Kirsh for insightful discussions and for providing constructive feedback on this manuscript.

## Authour contributions

S.M. performed ArsB expression and purification, cryo-EM sample preparation, data collection, processing and analysis; K.D. performed and analyzed arsenite resistance growth assays; S.M., W.M.C. and D.C.R. analyzed the data and wrote the manuscript; W.M.C. and D.C.R. supervised the project. All authors participated in revising the manuscript.

## Declaration of interests

The authors declare no conflicting interests.

## Notes

### Competing Interest Statement

The authors have declared no competing interest.

