## supplementary information for "Structural basis of metalloid transport by the arsenite efflux pump ArsB"

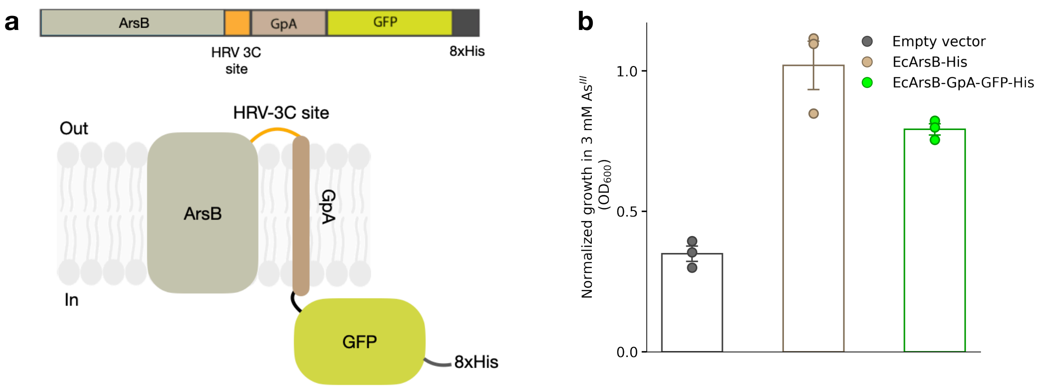


**Figure 1. *Lf*ArsB expression construct with GFP fusion. a** Schematic representation of the *Lf*ArsB expression construct bearing a C-terminal fusion of HRV-3C protease cleavage site – glycophorin A (GpA) – GFP – 8xHis. **b** Normalized growths of *E. coli* AW3110 cells bearing *Ec*ArsB-His or EcArsB-GFP fusion construct (*Ec*ArsB-GpA-GFP-His) as described in **a**, in presence of 3 mM $As^{III}$. $OD_{600}$ values in the presence of $As^{III}$ are normalized by corresponding values in the absence of $As^{III}$. Biological triplicates are reported, and error bars represent standard error of mean.

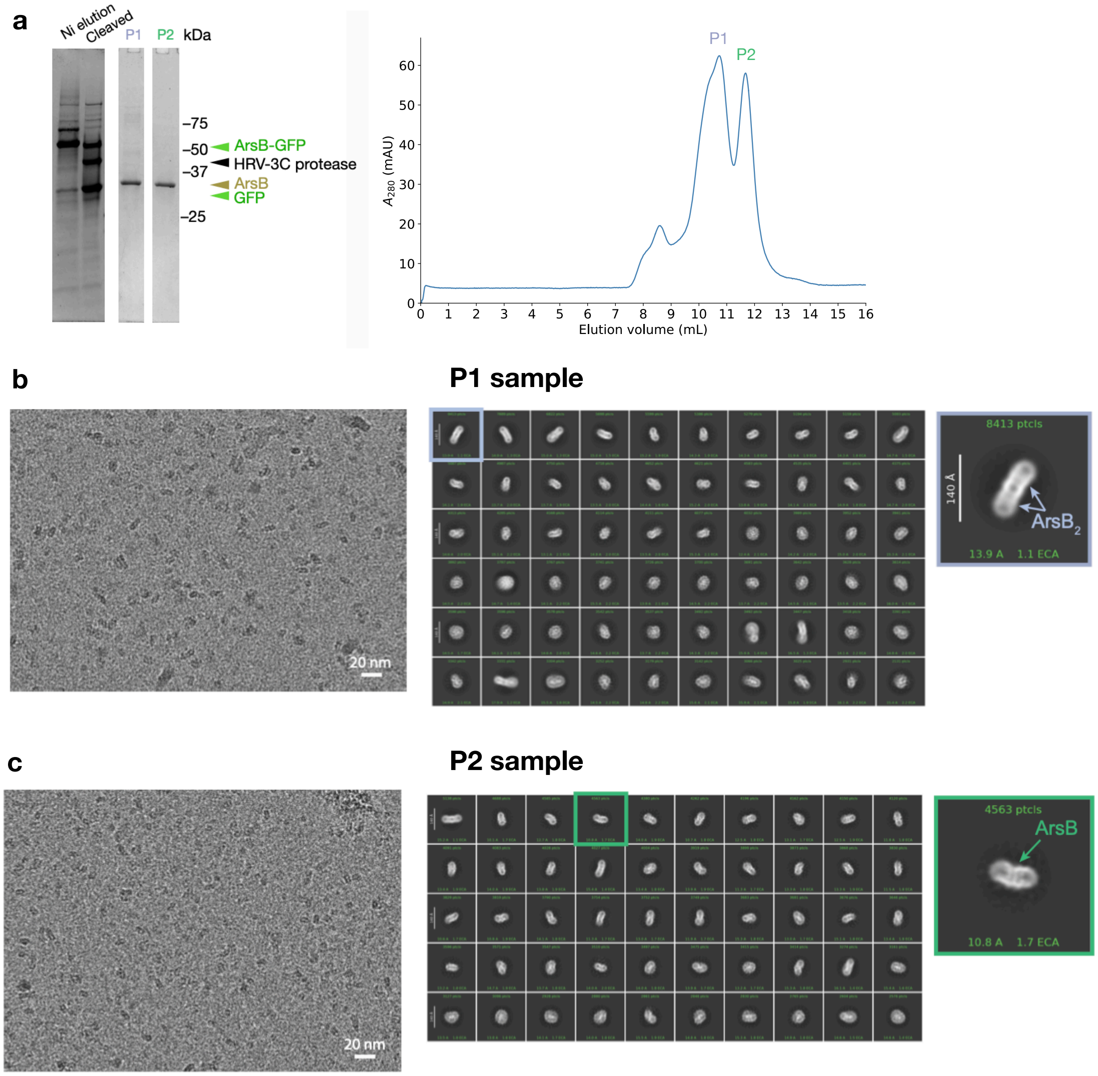


**Figure 2 . Purification and preliminary cryo-EM analysis of *Lf*ArsB solubilized in 0.03% (w/v) DDM. a** SDS-PAGE analysis (left) and SEC profile (right). The P1 and P2 samples, which show a pure ArsB band on the gel, correspond to peaks P1 and P2 on the SEC trace. Representative micrograph at 130,000x magnification and 2D class averages corresponding to P1 and P2 samples are shown in panels **b** and **c**, respectively. P1 sample is composed of ArsB dimer ($ArsB_2$) in a micelle, whereas the P2 sample is composed of ArsB monomer in a micelle.

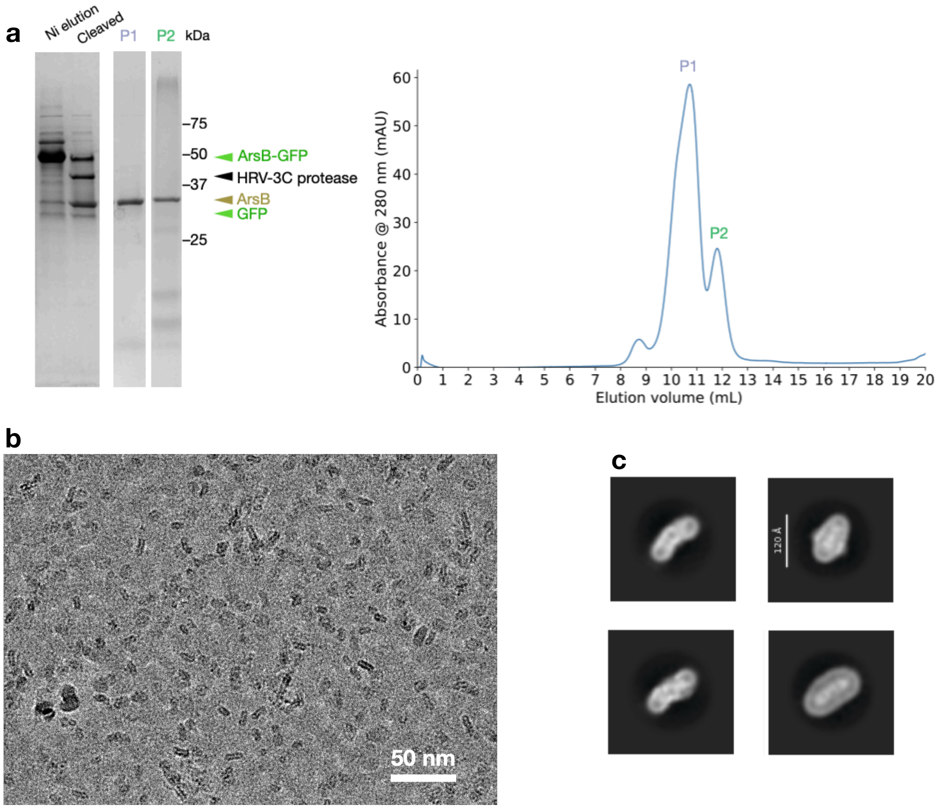


**Figure 3. Purification and preliminary cryo-EM analysis of *Lf*ArsB solubilized in 0.005%/0.0005% (w/v) LMNG/CHS. a** SDS-PAGE analysis (left) and SEC profile (right). **b** Representative micrograph from P1 sample at 130,000x magnification. **c** Representative 2D class averages from P1 sample showing ArsB dimers in micelles.

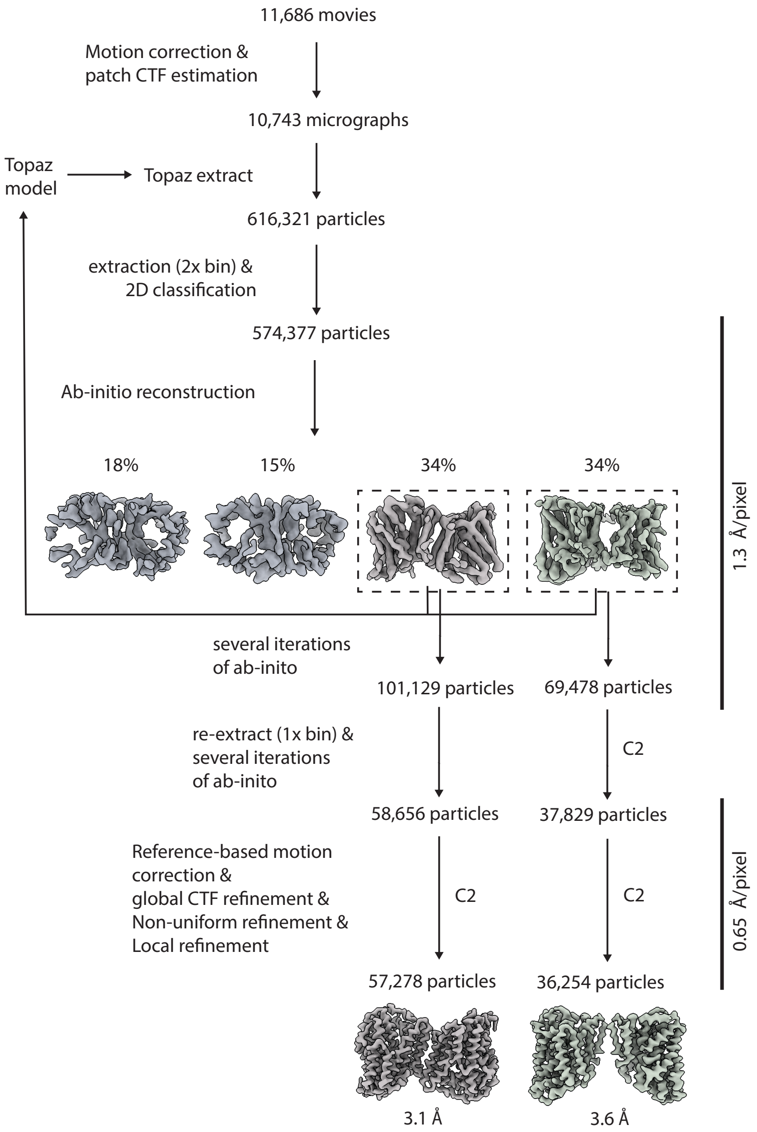


**Figure 4.** Cryo-EM data processing workflow for apo *Lf*ArsB structures in cryoSPARC.

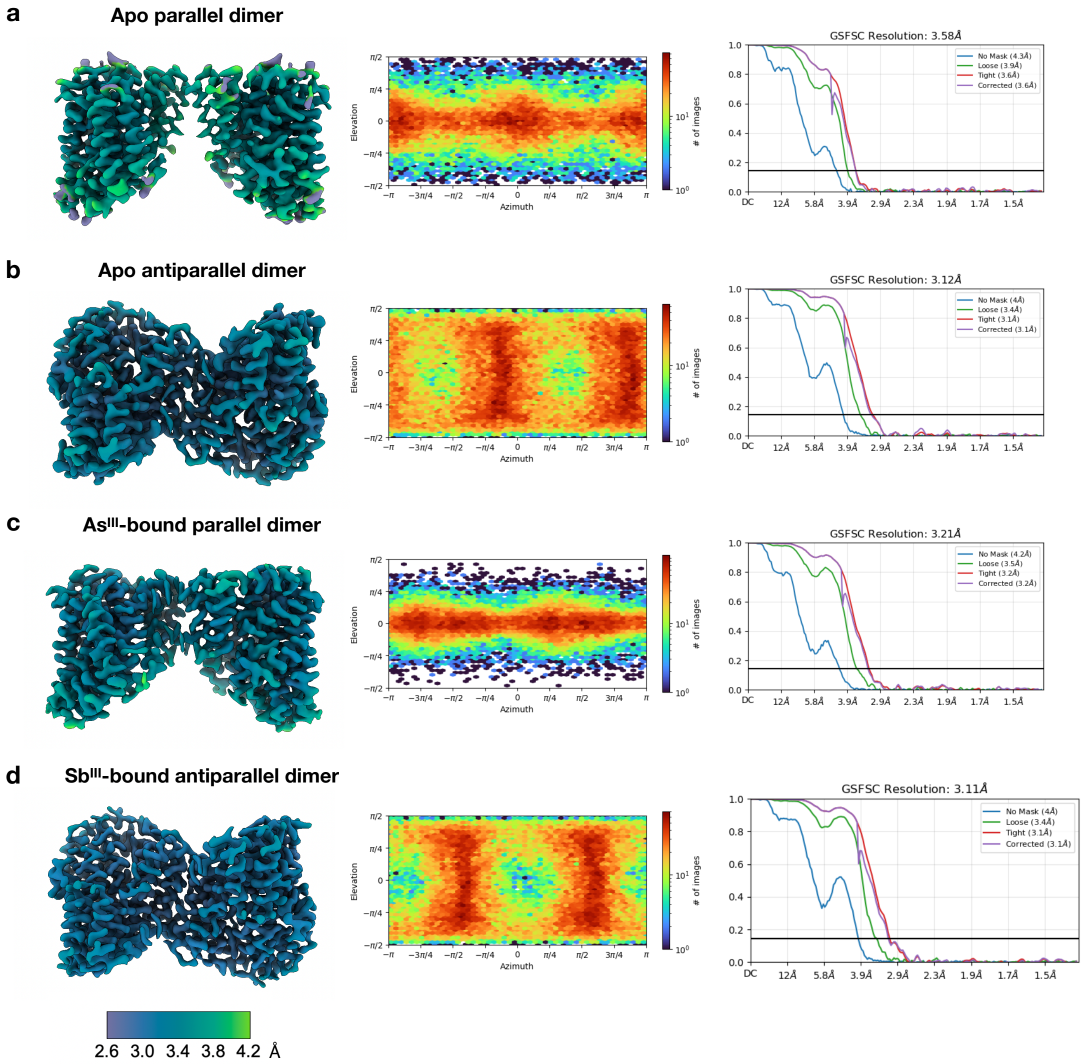


**Figure 5.** Cryo-EM maps and validation for **a** apo *Lf*ArsB, **b** *Lf*ArsB + $As^{III}$, and **c** *Lf*ArsB + $Sb^{III}$. Left, B-factor sharpened map colored by local resolution; center, angular distribution heatmap plot; and right, gold-standard Fourier shell correlation (GSFSC) curve (cut-off: 0.143).

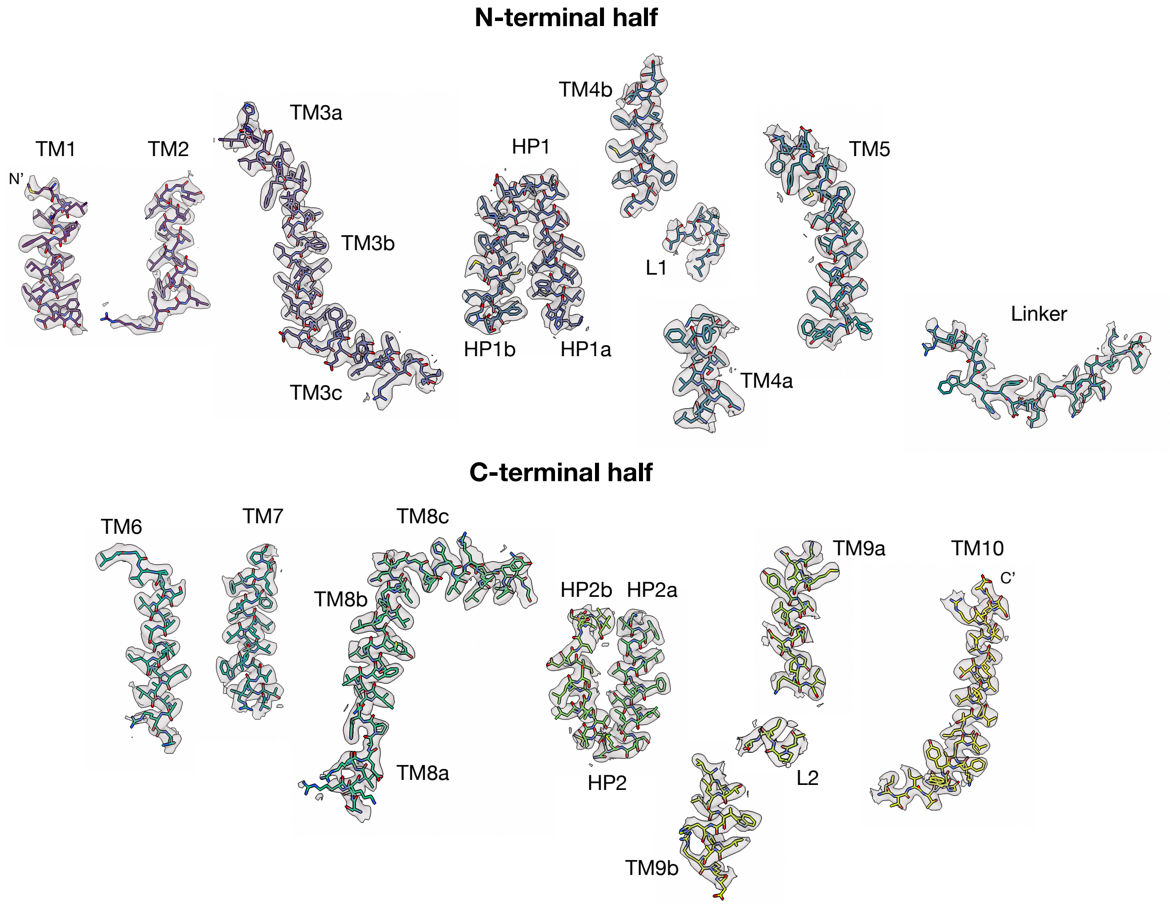


**Figure 6. Representative cryo-EM density maps for various segments of the apo antiparallel *Lf*ArsB structure.** Model is shown as sticks and colored using the viridis palette.

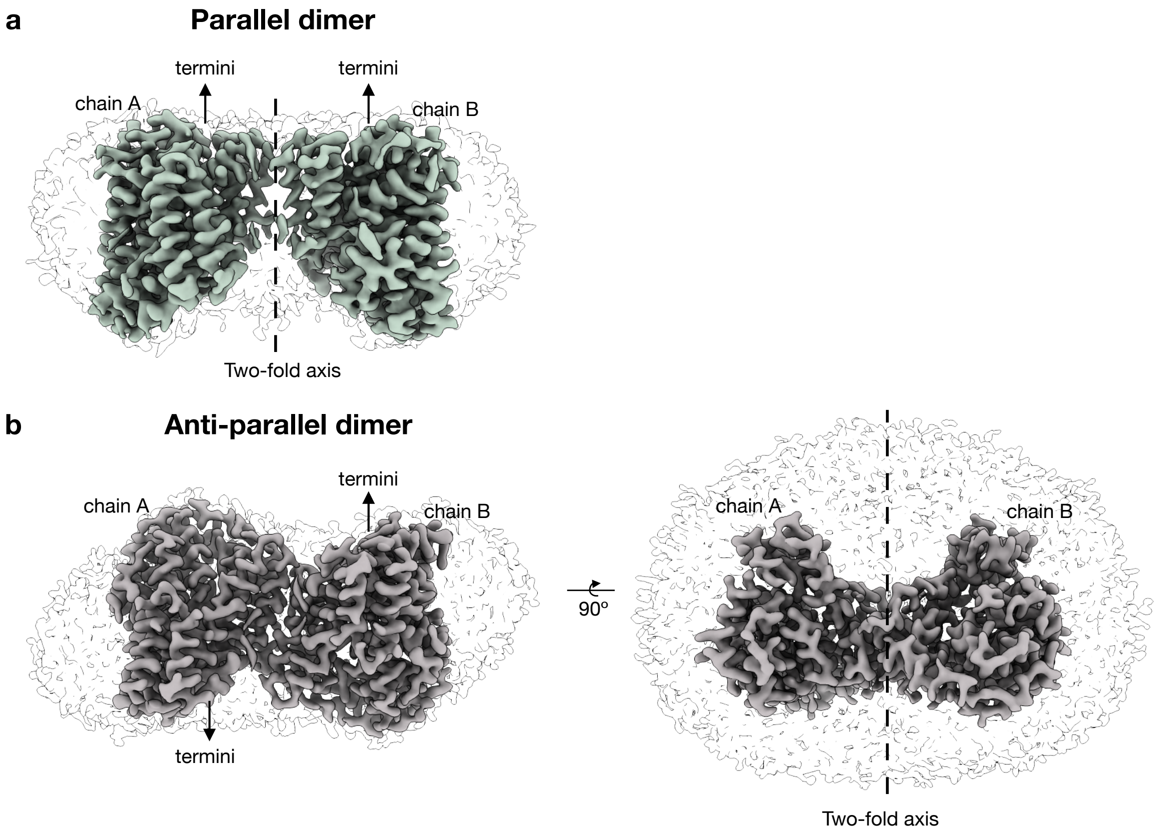


**Figure 7. Apo *Lf*ArsB dimer architectures in detergent micelles showing positions of the termini and the two-fold axis. a** Parallel dimer. **b** Antiparallel dimer.

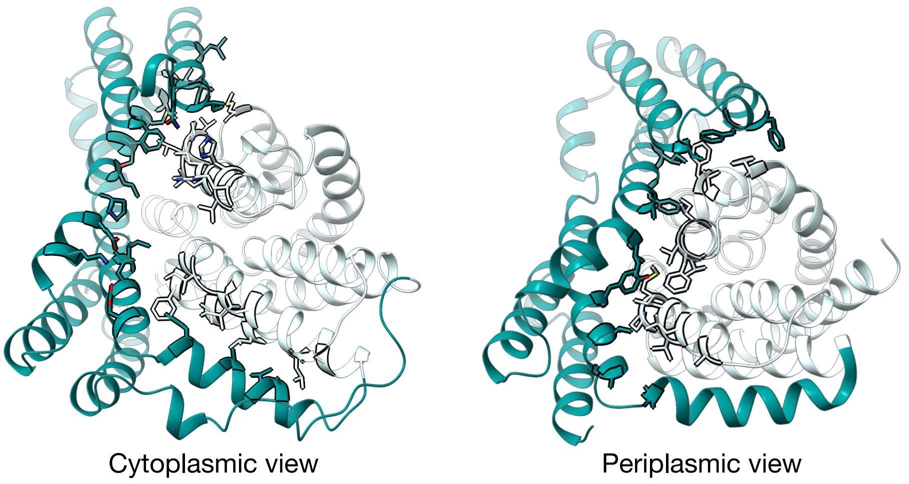


**Figure 8.** Residues (shown as sticks) at the interface of transport domain (cyan) and scaffold domain (teal) of *Lf*ArsB in the cytoplasmic view (left) and periplasmic view (right).

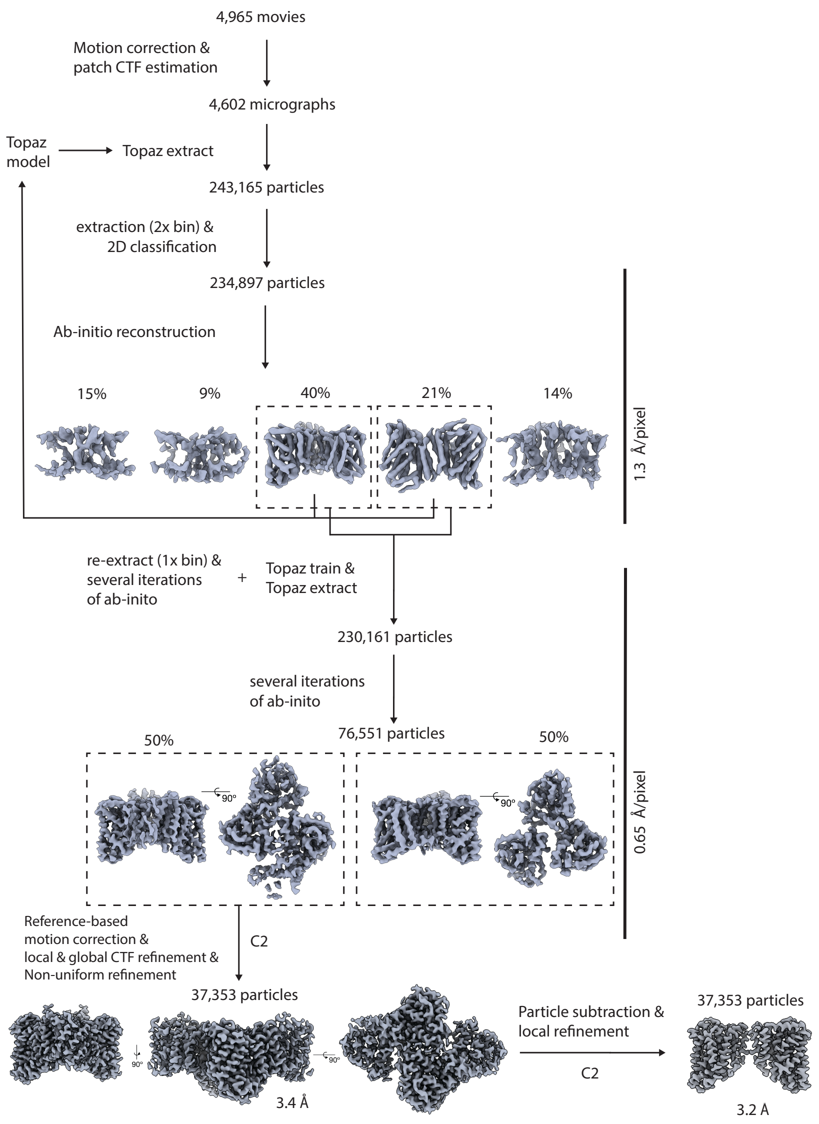


**Figure 9.** Cryo-EM data processing workflow for $As^{III}$-bound *Lf*ArsB structure in cryoSPARC.

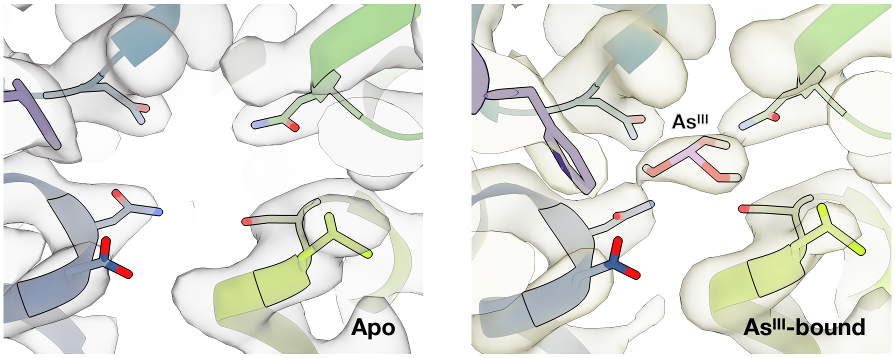


**Figure 10.** Metalloid-binding site of *Lf*ArsB apo structure (left) and As$^{III}$-bound structure (right) and corresponding Coulomb potential maps normalized and contoured at a threshold level of 7.0.

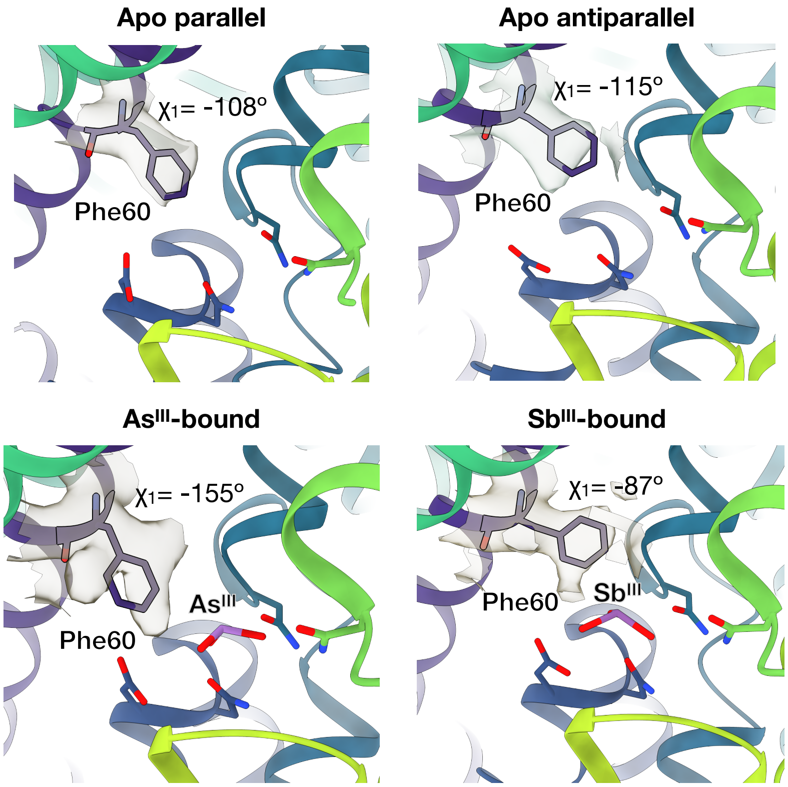


**Figure 11.** Multiple conformations of Phe60 sidechain across apo and metalloid-bound *Lf*ArsB structures. Respective $\chi_1$ torsion angles are labeled.

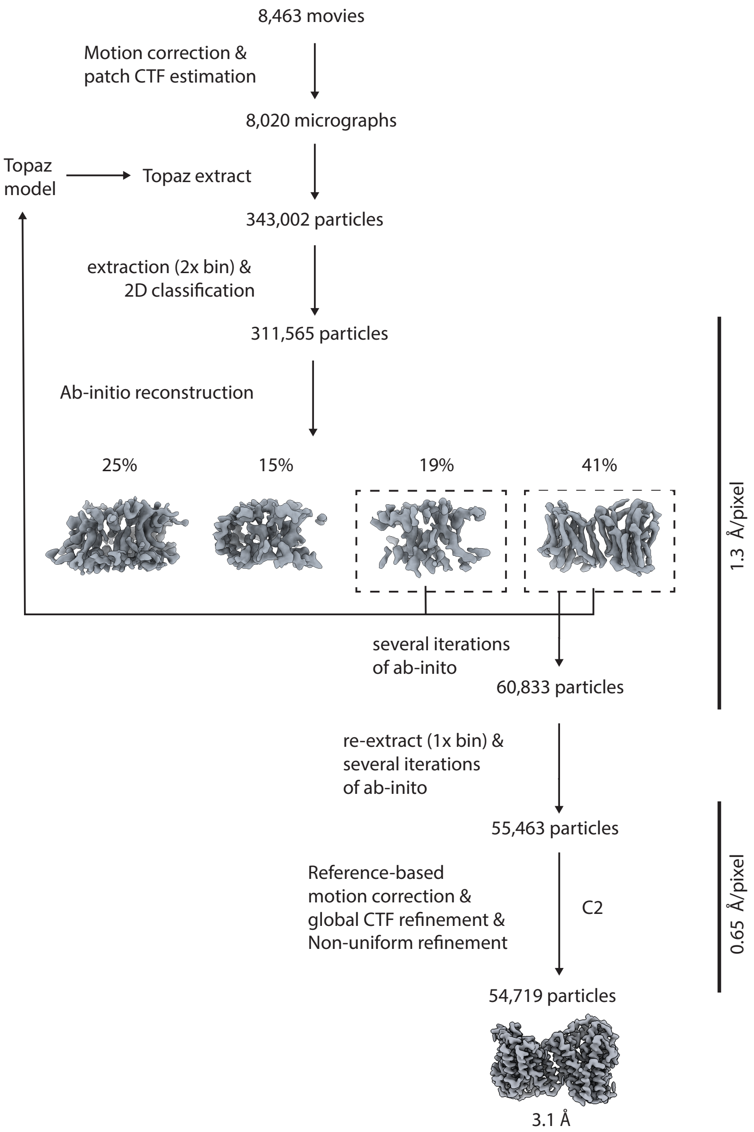


**Figure 12.** Cryo-EM data processing workflow for $Sb^{III}$-bound *Lf*ArsB structure in cryoSPARC.

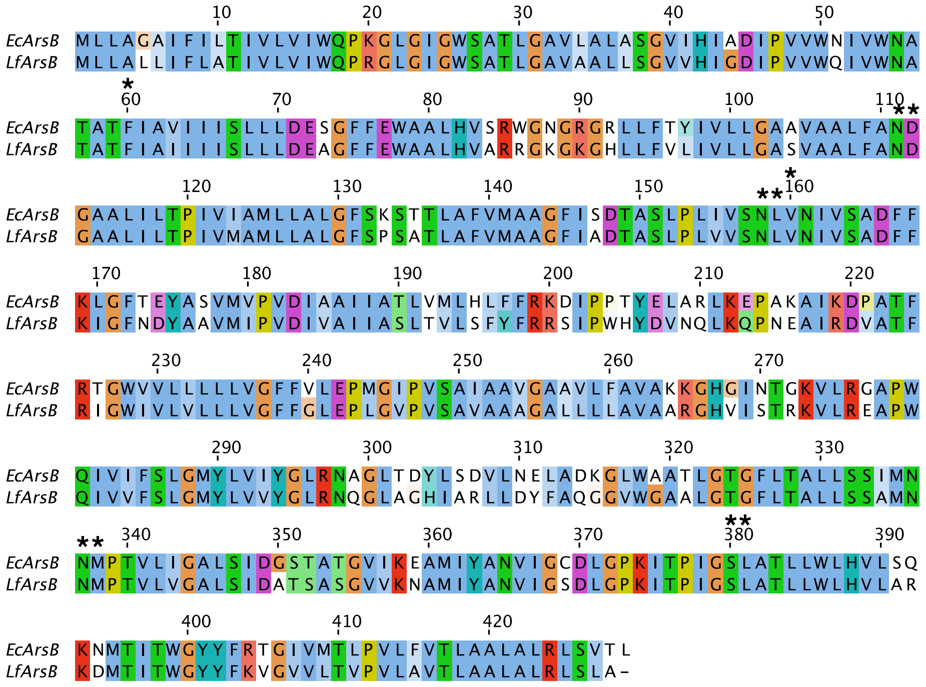


**Figure 13.** Sequence alignment of *Ec*ArsB and *Lf*ArsB prepared and visualized using Jalview[1]. Conserved residues of the metalloid-binding pocket are indicated with an asterisk (*) above the residue.

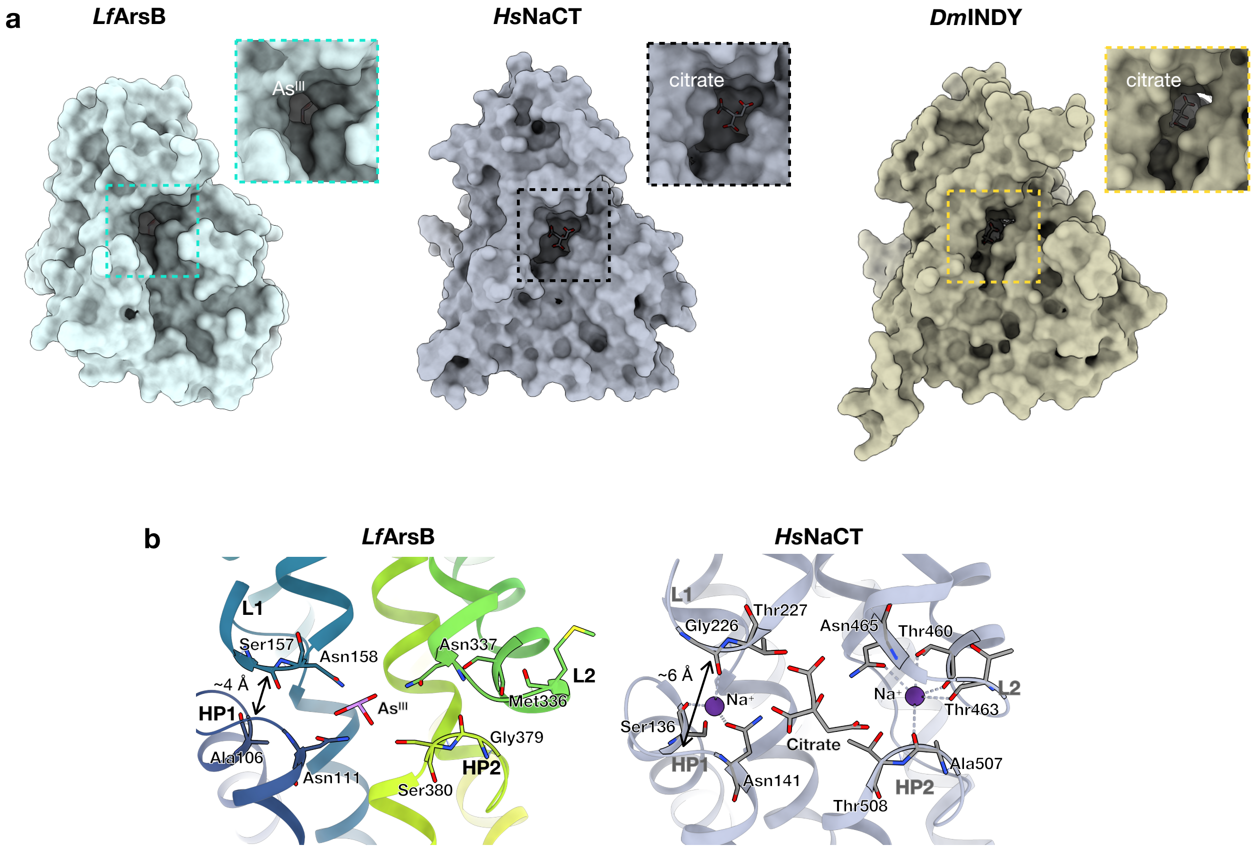


**Figure 14. Comparison of substrate-binding pockets of *Lf*ArsB (inward), *Hs*NaCT (inward-open), and *Dm*INDY (inward-occluded). a** Surface representation in cytoplasmic view highlighting the size of the substrate-binding pocket in each structure. **b** Substrate interaction residues and helix-loop motif residues for $Na^+$ interaction in *Lf*ArsB and *Hs*NaCT.

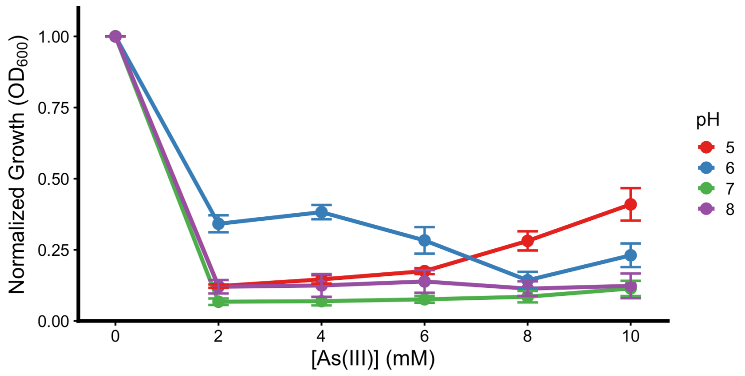


**Figure 15.** External pH-dependence of $As^{III}$ resistance conferred by empty pRSFdelT7 vector in AW3110 cells.

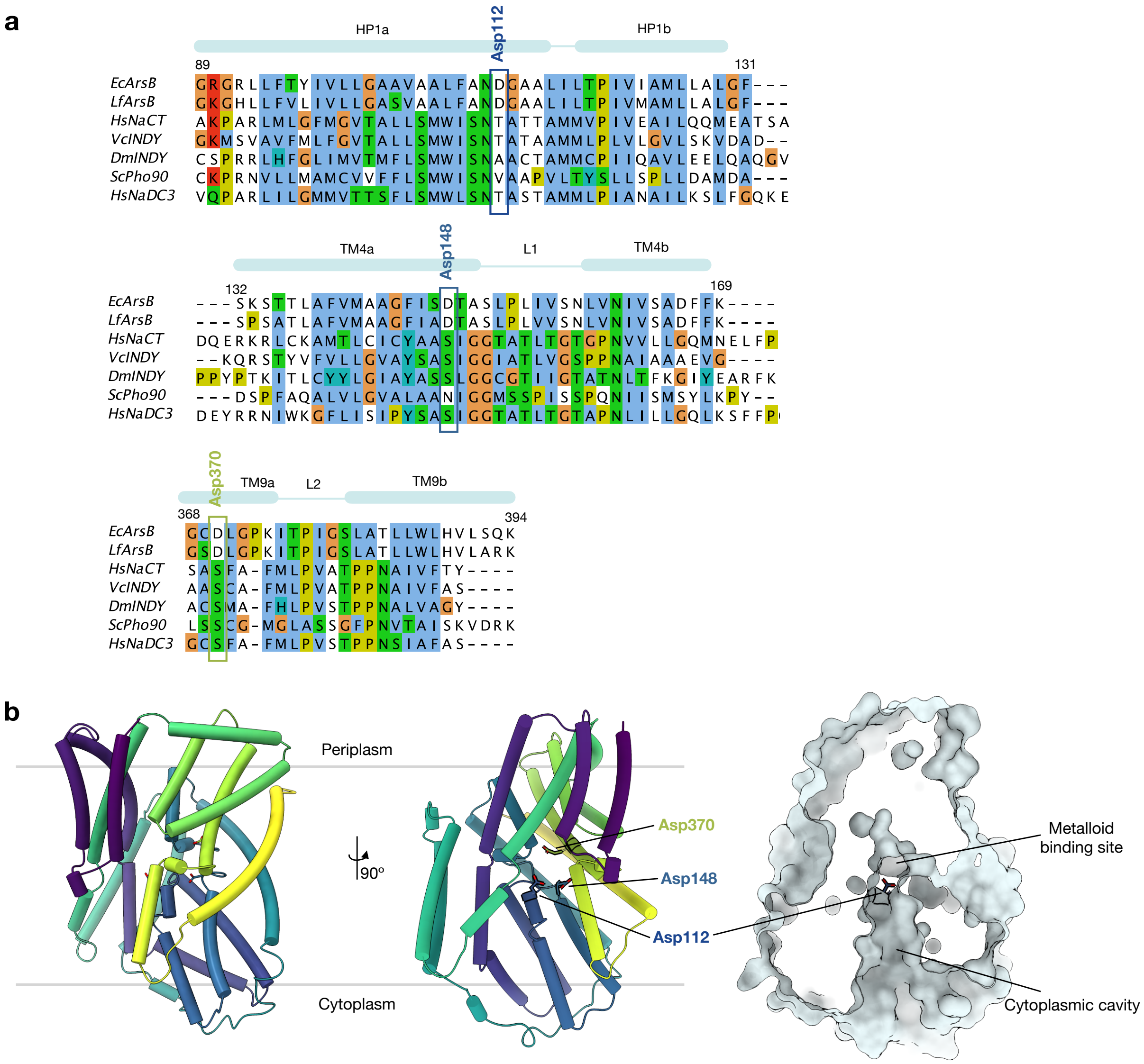


**Figure 16. H$^+$-coupling mechanism of *Lf*ArsB. a** Sequence alignment of ArsB and representative DASS transporters, highlighting that the putative H$^+$-coupling Asp residues are not found beyond ArsB sequences. Alignment was prepared using structure-based sequence alignment (Promals3D[2]) and visualized in Jalview[1]. Representative sequence limits shown correspond to ArsB sequences. **b** Positions of H$^+$-coupling Asp residues (Asp112, Asp148 and Asp370) in the *Lf*ArsB 'inward-facing' model.

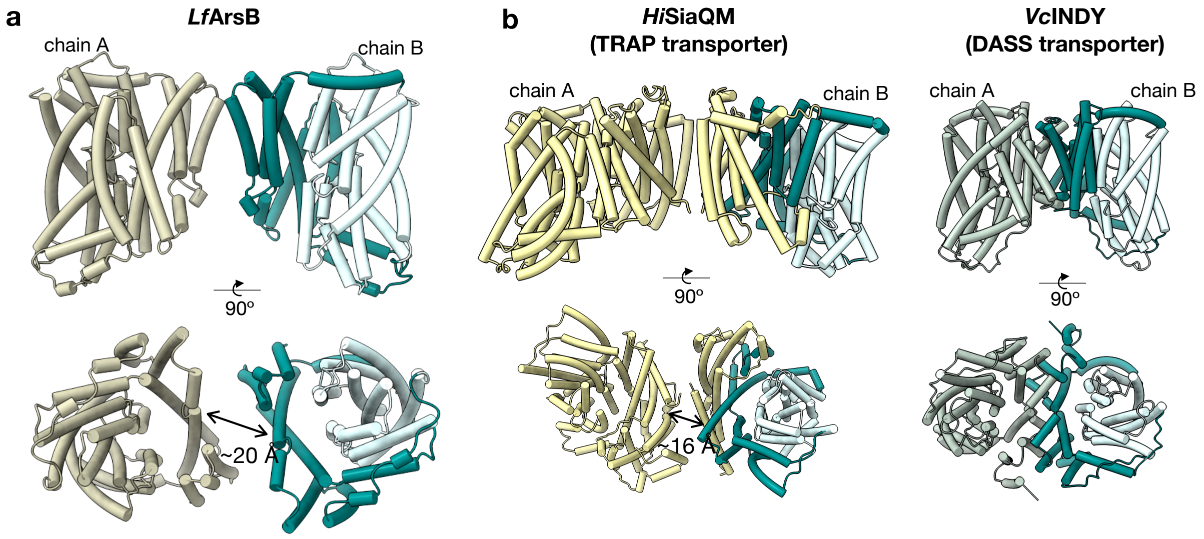


**Figure 17.** Parallel dimers of **a** *Lf*ArsB, and **b** TRAP transporter, SiaQM, from *H. influenzae* (*Hi*SiaQM; PDB 8THI) and DASS transporter, *Vc*INDY (PDB 7T9F), in two orientations.

**Table 1. Cryo-EM data collection, refinement and validation statistics.**

| | *Lf*ArsB apo parallel dimer (PDB ID 10TP) | *Lf*ArsB apo antiparallel dimer (PDB ID 10TQ) | *Lf*ArsB + $As^{III}$ parallel dimer (PDB ID 10TU) | *Lf*ArsB + $Sb^{III}$ antiparallel dimer (PDB ID 10UA) |
|---|---|---|---|---|
| **Data collection and processing** | | | | |
| Magnification | 130,000 | 130,000 | 130,000 | 130,000 |
| Voltage (kV) | 300 | 300 | 300 | 300 |
| Electron exposure ($e^-/Å^2$) | 70 | 70 | 70 | 70 |
| Defocus range (µm) | -0.8 to -2.8 | -0.8 to -2.8 | -0.8 to -2.8 | -0.8 to -2.8 |
| Pixel size (Å) (Super-resolution mode) | 0.325 | 0.325 | 0.325 | 0.325 |
| Movies | 11,686 | 11,686 | 4,965 | 8,463 |
| Total extracted particles | 616,321 | 616,321 | 316,729 | 343,002 |
| Final particles | 57,278 | 36,254 | 37,353 | 54,719 |
| Symmetry imposed | C2 | C2 | C2 | C2 |
| Map resolution (Å) (FSC 0.143 cut-off) | 3.6 | 3.1 | 3.2 | 3.1 |
| **Refinement** | | | | |
| Initial model used | *Lf*ArsB AlphaFold model | *Lf*ArsB AlphaFold model | *Lf*ArsB AlphaFold model | *Lf*ArsB AlphaFold model |
| Model composition: | | | | |
| Protein residues | 856 | 856 | 856 | 856 |
| Ligands | | | $As(OH)_3$ | $Sb(OH)_3$ |
| Map sharpening *B* factor ($Å^2$) | -117 | -112 | -92 | -119 |
| R.m.s. deviations: | | | | |
| Bond lengths (Å) | 0.007 | 0.008 | 0.008 | 0.007 |
| Bond angles (°) | 0.687 | 0.703 | 0.603 | 0.587 |
| **Validation** | | | | |
| MolProbity score | 1.30 | 1.30 | 1.46 | 1.54 |
| Clashscore | 5.61 | 5.53 | 5.68 | 5.68 |
| Poor rotamers (%) | 0 | 0 | 0 | 0 |
| Ramachandran plot: | | | | |
| Favored (%) | 98.4 | 98.4 | 97.2 | 96.5 |
| Allowed (%) | 1.6 | 1.6 | 2.8 | 3.5 |
| Outliers (%) | 0 | 0 | 0 | 0 |
